# A spring-loaded and leakage-tolerant synthetic gene switch for in-vitro detection of DNA and RNA

**DOI:** 10.1101/2024.02.12.579921

**Authors:** Krishna Gupta, Elisha Krieg

**Affiliations:** Institute for Biofunctional Polymer Materials, Leibniz Institute of Polymer Research Dresden, Germany; Faculty of Chemistry and Food Chemistry, Technische Universität Dresden, Germany

**Keywords:** DNA Nanotechnology, Synthetic Biology, Diagnostics, Point-of-Care, Strand Displacement

## Abstract

Nucleic acid tests (NATs) are essential for biomedical diagnostics. Traditional NATs, often complex and expensive, have prompted the exploration of Toehold-Mediated Strand Displacement (TMSD) circuits as an economical alternative. However, the wide application of TMSD-based reactions is limited by ‘leakage’—the spurious activation of the reaction leading to high background signals and false positives. Here we introduce a new TMSD cascade that recognizes a custom nucleic acid input and generates an amplified output. The system is based on a pair of thermodynamically spring-loaded DNA modules. The binding of a predefined nucleic acid target triggers an intermolecular reaction that activates a T7 promoter, leading to the perpetual transcription of a fluorescent aptamer that can be detected by a smartphone camera. The system is designed to permit the selective depletion of leakage byproducts to achieve high sensitivity and zero-background signal in the absence of the correct trigger. Using Zika virus (ZIKV)- and severe acute respiratory syndrome coronavirus 2 (SARS-CoV-2)-derived nucleic acid sequences, we show that the assay generates a reliable target-specific readout. Native RNA can be directly detected under isothermal conditions, without requiring reverse transcription, with a sensitivity as low as 200 attomole. The modularity of the assay allows easy re-programming for the detection of other targets by exchanging a single sequence domain. This work provides a low-complexity and high-fidelity synthetic biology tool for point-of-care diagnostics and for the construction of more complex biomolecular computations.

## INTRODUCTION

Nucleic acid tests (NATs) are important for the monitoring of infectious diseases. The detection of pathogen-specific DNA and RNA is the precondition for timely isolation, treatment, and disease mapping. Consequently, NATs have played a crucial role in fighting the COVID-19 pandemic. Laboratory-based tests, such as reverse transcription-quantitative polymerase chain reaction (RT-qPCR), are considered the gold standard for identifying infectious diseases^1^. They typically involve several enzymatic reactions, first reverse-transcribing RNA to DNA, followed by multiple cycles of thermal denaturation and step-wise amplification. Despite their excellent sensitivity and reliability, their high complexity, resource requirements, bulky setup, and processing time make them poorly suitable for point-of-care (POC) diagnostic testing. Notably, the global usage of RT-qPCR during the COVID-19 pandemic caused a bottleneck of reagents, test kits, and enzymes, affecting the control and monitoring of disease propagation.

There is hence a large interest in simpler and more accessible technologies that are suitable for POC diagnostics. An ideal POC test operates under isothermal conditions, thus obviating the need for thermocycling instruments. Loop-Mediated Isothermal Amplification (LAMP) is amongst the most popular isothermal tests^2^. Other important isothermal methods are Nucleic Acid Sequence-based Amplification (NASBA)^3^, Recombinase Polymerase Amplification (RPA)^4^, and, more recently, techniques based on the CRISPR/Cas enzyme system^5–9^. Yet, all of these methods require complex mixtures of reagents and enzymes (in particular for the detection of RNA). The reagents are typically expensive, prone to supply shortages, require cold storage, and are unsuitable for use in resource-limited areas. Current isothermal amplification techniques are also susceptible to false positives due to mispriming, thus requiring extensive primer screening and trial-and-error testing^10–13^. Therefore, there remains an unmet need for fast, programmable, low-cost, and low-complexity assays for the detection of infectious diseases^1,14^.

Toehold-mediated strand displacement (TMSD) cascades offer a promising potential solution to this challenge^15–21^. TMSD involves the binding of an *input* DNA molecule to the *toehold* (a single-stranded overhang domain) of a self-assembled DNA construct. Once bound, the input displaces and thereby activates an *output* strand. This simple signal translation mechanism can be used to build complex reaction networks that operate without enzymatic assistance. Catalytic hairpin assembly (CHA)^22^ and hybridization-chain reaction (HCR)^23–25^ are amongst the most explored TMSD-based assays for enzyme-free biosensing and diagnostics. TMSD-based assays usually require meticulous sequence screening and the use of expensive fluorescently labeled strands to generate a detectable output^26–28^. Moreover, TMSD reactions are plagued by *leakage*^26,29,30^— false positive results due to undesirable crosstalk and inadvertent signal activation—though significant progress is being made towards low-leakage and high-fidelity system designs^28,31–34^.

Researchers have also begun combining TMSD reactions with enzymes. Notably, transcriptional switches such as *genelet* circuits^35–37^, toehold switches^38^, and systems involving allosteric transcription factors^39^ have been developed to regulate RNA or protein expression in vitro. These approaches hold great promise to directly process information within a biological specimen, and subsequently generate a refined diagnostic output.

Inspired by these innovations, and motivated by recent disease outbreaks, we aimed to develop a detector for pathogen-specific nucleic acids. The detector design combines four key features: (1) simplicity: comprising a small number of rapidly accessible components and no more than one enzyme; (2) versatility: detection of arbitrary nucleic acid sequences—both single stranded DNA (ssDNA) *and* RNA (ssRNA); and (3) modularity: the system is easily re-programmable for quick adaptation to future pathogen outbreaks; (4) robustness: the system’s response is highly specific to the selected target sequence, exhibiting negligible interference from background sensor leakage.

## MATERIALS AND METHODS

### Solvents and reagents

All solvents and reagents were purchased from commercial vendors and used as received unless otherwise specified. Molecular biology grade water was obtained from a Milli-Q system. Ammonium persulfate (Cat. #A3678), Tris-EDTA buffer (Cat. # T9285), 3,5-difluoro-4-hydroxy-benzylidene imidazolinone (DFHBI, Cat. # SML1627), Potassium Chloride 3M (Cat. # 60137) from Sigma-Aldrich. Molecular biology grade tetramethylethylenediamine (TEMED, Cat. #15524010), SYBR™ Gold (Cat. #S11494), TBE buffer (Cat. # 15581044), DNA ladders (Cat. # SM1211 & # SM0311), DNA loading dye (Cat. # R0611) from Thermo Fischer Scientific. ATP (Cat. # NU-1010), GTP (Cat. # NU-1012), CTP (Cat. # NU-1011), and UTP (Cat. # NU-1013) were purchased from Jena Bioscience. RNA loading dye (Cat. # B0363S), T7 RNA polymerase (Cat. # M0251), ssRNA ladder (Cat. # N0364), recombinant albumin (Cat. # B9200), and dNTPs solution mix (Cat. # N0447) from New England Biolabs (NEB). Nuclease free water (Merck, Cat. # W4502), and Molecular Biology grade 19:1 acrylamide/bis-acrylamide (Serva Electrophoresis, Cat. #10679.01) was procured from other commercial vendors. All DNA oligonucleotides were purchased from Integrated DNA Technologies (IDT) as desalted oligos, except P and S strands, which were PAGE-purified Ultramers™. Synthetic DNA stock solutions were stored at 100 μM concentration in 1X IDTE buffer (IDT, Cat. #11-05-01-09, pH 8.0). Cytoplasmic RNA from HEK293T cells was provided by the lab of Prof. Dr. Sun Hur (Harvard Medical School)^40^. A purified RNA calibration standard from cultured SARS-CoV-2 (B.1.1.7; 3.96x10^8^ cp/μL) was obtained from the Institute for Medical Microbiology and Virology, University Clinic Carl Gustav Carus, Dresden, Germany. The HPLC-purified RNA oligonucleotides were ordered from IDT and biomers.net GmbH, dissolved in nuclease-free water (100 μM stock concentration), and stored as 1-5 μl aliquots at -80 °C.

### Nucleic acid design and analysis

The oligonucleotides were designed and predicted structures were analyzed using the NUPACK webserver (https://nupack.org/)^41^, IDT’s oligo analyzer (https://eu.idtdna.com/calc/analyzer), and the OligoCalc webserver (http://biotools.nubic.northwestern.edu/OligoCalc.html)^42^. TMSD cascade sequences were designed using the NUPACK design tool. Specific design parameters are described in Supplementary Notes 2.1 and 2.2. The Zika virus binding sequence was selected from a previously reported recombinase polymerase amplification (RPA) reverse primer^43^, which was trimmed to 33 nt length (Figure 4a). The detailed selection principles for the pathogen-specific binding sequence is explained in Supplementary Note 2.3. The selected target-recognition sequences (Figure 4a) were directly used for the assay without requiring further experimental screening. All sequences of oligonucleotides used in this study are listed in Supplementary Table S2.

### Polyacrylamide gel electrophoresis (PAGE)

All PAGE gels were cast on XCell SureLock Mini-Cell® Electrophoresis system (Invitrogen, Cat. #SM1211) using Bolt™ Mini Gel Cassettes (Invitrogen, Cat. # NW2010). Native polyacrylamide gels of different percentages were prepared using a 40% (w/w) 19:1 acrylamide/bis-acrylamide in 0.5X Tris-Borate-EDTA buffer. The samples were run at 110-170 V, using a Consort EV265 power supply, stained using 1X SYBR™ Gold, and imaged on a Typhoon FLA 9500 laser scanner (GE Healthcare Life Sciences) using their software using a blue LD laser (excitation at 473 nm), BPB filter, 400 Volts setting, and with the resolution of 10-50 μm/pixel. The image analysis and densitometric quantification of gel bands was performed using ImageJ (v. 1.52a)^44^.

### DNA construct assembly

Constructs **[P]** and **[S]** were assembled by combining base strands with different equivalents of blocking strands (cf. Table 1 and Figure 2a) in an annealing buffer (40 mM Tris-HCl, 300 mM KCl, pH 7.9). The samples were split into PCR tubes with 50 μl volume each. The samples were annealed by heating to 80 °C for 1 min, followed by a temperature ramp from 70 °C to 20 °C at the rate of -0.5 °C min^-1^. The assembled constructs were stored at 4 °C for a minimum of 8 hours before use.

**Table 1.** Typical concentrations of base strands (P or S) and the amount of blocking strands in equivalents (eq.) relative to the base strand. Design changes in the cascade versions are described in Supplementary Note 2.2 and Supplementary Figure S5.

| Cascade version | [Base strand] | Blocking strands |
| --- | --- | --- |
| 1 | 200-500 nM | 2 or 3 eq. |
| 2 (leakage-reduced) | 200 nM | 10 eq. |
| 3 (leakage-tolerant) | 200 nM | 3 eq. |

### SARS-CoV-2 mimic RNA expression and purification

The transcription template for the SARS-CoV-2-specific sequence was produced by PCR amplification of the N-gene Positive Control plasmid (IDT, #10006625) as a template, using the NEBNext Ultra II Q5 Master Mix (NEB, #M0544), primers (0.5 μM each, Supplementary table S3) using the qPCR program in Supplementary Table S4. The forward primer contained a T7 adapter. The PCR product was purified using the DNA Clean and Concentrator-5 kit (Zymo Research, #D4004) and transcribed using the HiScribe T7 High Yield RNA Synthesis Kit (NEB, Cat. # E2040S), template concentration 10 nM, 37 °C for 16 hours. After DNAse I treatment (NEB, Cat. #M0303), the RNA was purified with the Monarch RNA Cleanup Kit (NEB, Cat. #T2050) and run on a 1.5% agarose gel with 0.3% hydrogen peroxide (Supplementary Figure S7a)^45^. Preliminary validation of concentration and integrity was done using Implen P360 Nanophotometer and Agilent Bioanalyzer 2100 (Supplementary Figure S7b). The RNA product was supplemented with RNAse inhibitor (NEB, #M0307), aliquoted, and stored at -80 °C. The RNA concentration was quantified using a SARS-CoV-2 triplex PCR kit (Astra Biotech, Cat. #89-03 Form F), and known concentrations of SARS-CoV-2 RNA sequences were used as calibration standards (Supplementary Figure S7c) using two independent experiments.

### Strand displacement assay

The strand displacement assays were conducted by adding the respective DNA constructs, triggers, and other oligos components to the reaction buffer (40 mM Tris–HCl pH 7.9, 20 mM MgCl_2_, and 50 mM KCl) unless specified otherwise. These experiments were conducted on a 10-20 μl scale. The samples were incubated at a constant temperature (25 °C or 37 °C) for the required time and were analyzed on a native PAGE gel.

### Nucleic acid detection using leakage-reduced constructs

The leakage-reduced constructs we assembled using sequences from TMSD Cascade 2_15nt_COVID and TMSD Cascade 2_15nt_ZIKV with 10 eq. of blocking strands. The assay was conducted at a 20 μl scale using two steps. Firstly, the strand displacement was conducted by incubating DNA constructs (20 nM each) with different concentrations of triggers in a reaction buffer (40 mM Tris–HCl pH 7.9, 20 mM MgCl_2_, 50 mM KCl, 10 mM dithiothreitol, 80 μM DFHBI, 5 mM each rNTP, and 0.5 mg/ml recombinant albumin) for 30 minutes. Different concentration of Zika virus (ZIKV_ssDNA & ZIKV_ssRNA) and SARS-CoV-2 (COVID_ssDNA, COVID_ssRNA & SARS-CoV-2 mimic) triggers were freshly prepared by serial dilution in DNA LoBind tubes (Eppendorf) using nuclease-free water. Cytoplasmic RNA (∼800 ng) was used as negative control, nuclease-free water as leakage reference, and **[P]** and **[S]** as internal references. Next, the samples were transferred to a white qPCR plate (Sarstedt, Cat. #72.1981.232), and T7 RNA polymerase (5 U/μl) was added to all samples, followed by incubation at 37 °C for up to 7 hours on a real-time qPCR instrument (Biorad, CFX96). To measure the time-dependent change in fluorescence, the instrument was set to carry out 1-minute plate reads at 20°C within 10-minute intervals.

### Nucleic acid detection using leakage-tolerant assay

To constructs were assembled at 200 nM base strand concentration in annealing buffer, using sequences from TMSD Cascade 2_15nt_COVID and TMSD Cascade 2_15nt_ZIKV and 3 eq. concentration of biotin-TEG capped blocking strands. The assay was conducted in three steps: strand displacement, leakage capture, and signal amplification at a 15μl scale. Firstly, the two constructs (1.5 μl each) were combined with the 1.5 μl trigger strand solution (varying trigger concentrations) and 4.2 μl strand displacement buffer (final concentrations: 40 mM Tris-HCl, pH 7.9, 12 mM MgCl_2_, 150 mM KCl). The mixture was incubated at 37 °C for 30 minutes. Different concentrations of triggers (ZIKV and SARS-CoV-2), negative control, and references (leakage, internal controls) were used as described before. Next, the resulting displacement product was added to tubes containing pre-washed streptavidin-coated magnetic beads Dynabeads™ MyOne™ Streptavidin C1 (Thermo Scientific, #65001). For each sample, 10 μl magnetic beads were washed by removing the storage buffer, adding a wash buffer (40 mM Tris-HCl, pH 7.9, 20 mM MgCl_2_, 50 mM KCl), briefly shaking the tube and removing the buffer, leaving magnetic beads in the tube. The samples were incubated with magnetic beads for 30 minutes with neoLabLine rotator-vortexer (neolab, Cat. #7-0045) using a homemade 3D printer shaker attachment and then captured using a homemade 3D printed magnetic separation stand. The supernatant was extracted and added to 6.3 μl expression mix (10 mM dithiothreitol, 80 μM DFHBI, 5 mM each NTP, 0.5 mg/ml recombinant albumin, 5 U/μl T7 RNA polymerase) on a white qPCR plate. Fluorescence was recorded as described before.

### Fluorescence quantification and statistical analysis

Baseline subtractions for data pre-processing were performed using sample containing **[S]** construct selected as the baseline sample. For curve fitting, the first fluorescence intensity value of each sample was subtracted. The slope was calculated after 250 minutes, and the signal-to-leakage value was calculated as:

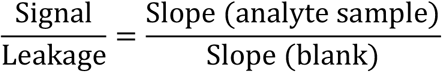

The signal-to-leakage vs concentration values were plotted on GraphPad Prism 7. Standard deviations, mean values, and regression plots were generated in the same software. L_c_ was determined by calculating the intersection point between the signal-to-leakage baseline and the linear regression for the target concentrations that generate signals at least one standard deviation above the baseline^46,47^.

## RESULTS

### The sensor is based on a thermodynamically spring-loaded pair of DNA modules

Here, we present a nucleic acid detection method that combines a TMSD cascade with in vitro transcription of a fluorescent light-up aptamer by a T7 RNA polymerase (Figure 1). The system contains a pathogen-recognizing DNA construct **[P]** and a second construct **[S]**, which encodes the Spinach^48,49^ aptamer. The modularity of the sensor offers easy adaptation to different target sequences by exchanging a single domain on **[P]**. The detection is carried out in a single tube containing at least 10 μL total reaction volume. In the presence of a predefined DNA or RNA target, the system generates a fluorescent signal under isothermal conditions. Unlike most TMSD assays, none of the DNA strands require labeling by a fluorophore or fluorescence quencher, as the fluorescent signal is generated via in-situ self-assembly of the expressed aptamer with a conditional fluorophore.

**Figure 1.**
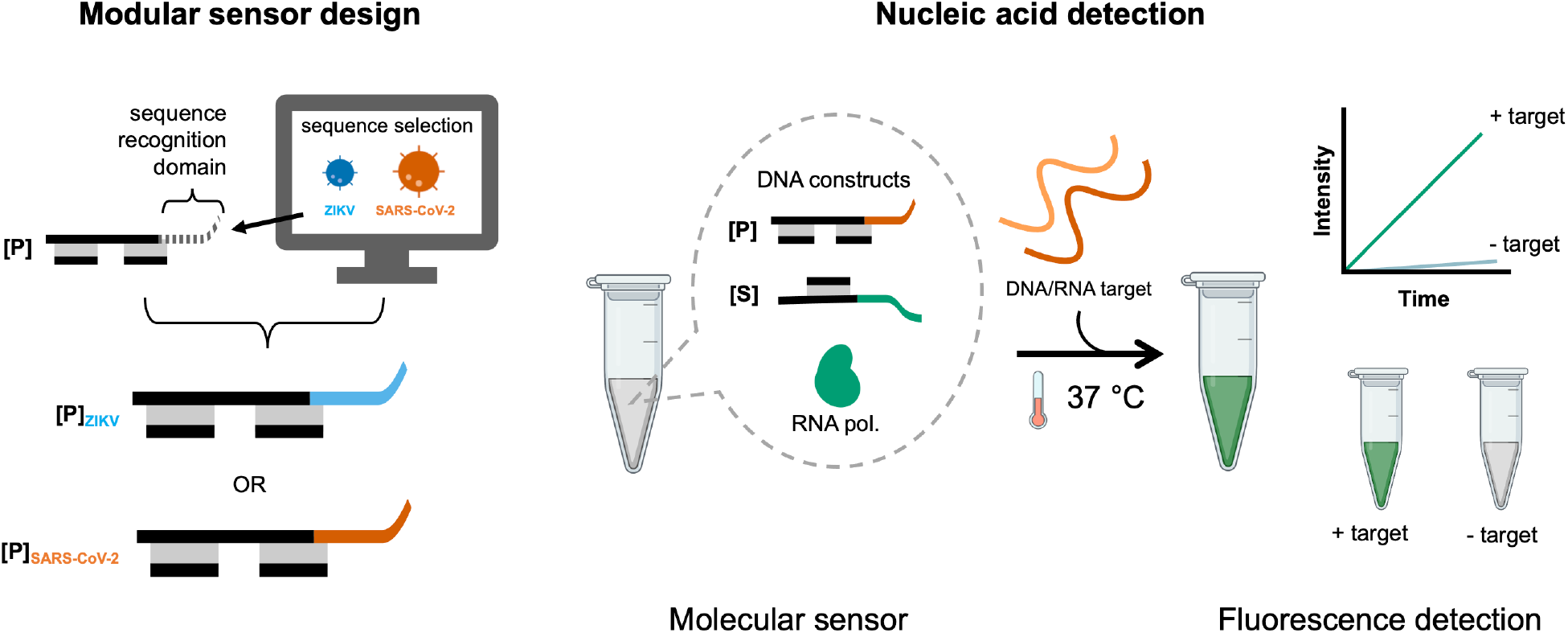
The modularity of the DNA-based sensor allows rapid assay development and quick adaptation to new pathogen sequences. The reaction mixture consists of two DNA constructs: a pathogen-interacting construct **[P]** and a Spinach aptamer-encoding construct **[S]**. The reaction is initiated by a predefined single-stranded nucleic acid target, triggering the transcription of a fluorescent Spinach aptamer in the presence of T7 RNA polymerase at 37 °C. A new nucleic acid detection assay can be developed by exchanging a single domain in the **[P]** construct.

The detailed sensor design is depicted in Figure 2a. **[P]** comprises a major strand (P) and two blocking strands (B_x_ and B_z_). P contains a T7 RNA polymerase promoter^50^ domain (*t7p*) and a pathogen-recognizing domain (*path*). **[S]** is assembled from a major strand (S) and one blocking strand (B_y_). S is largely complementary to P, but instead of *path*, it contains a template sequence for the Spinach aptamer (*spin*). There is a large thermodynamic driving force for the hybridization reaction **[P]** + **[S] → [PS]** + B_x_ + B_y_ + B_z_, where **[PS]** is the P + S hybridization product (ΔG ≤ -400 kJ mol^-1^, see Supplementary Notes 2 and Supplementary Table S1). However, the blocking strands prevent the initiation of this exchange reaction in the absence of a trigger. An equimolar **[P]** + **[S]** mixture is therefore trapped in a spring-loaded dormant state that can be stable for days. Importantly, the polymerase-binding part of the *t7p* domain in **[S]** is largely single stranded, rendering it incapable of recruiting T7 polymerase^51^.

**Figure 2.**
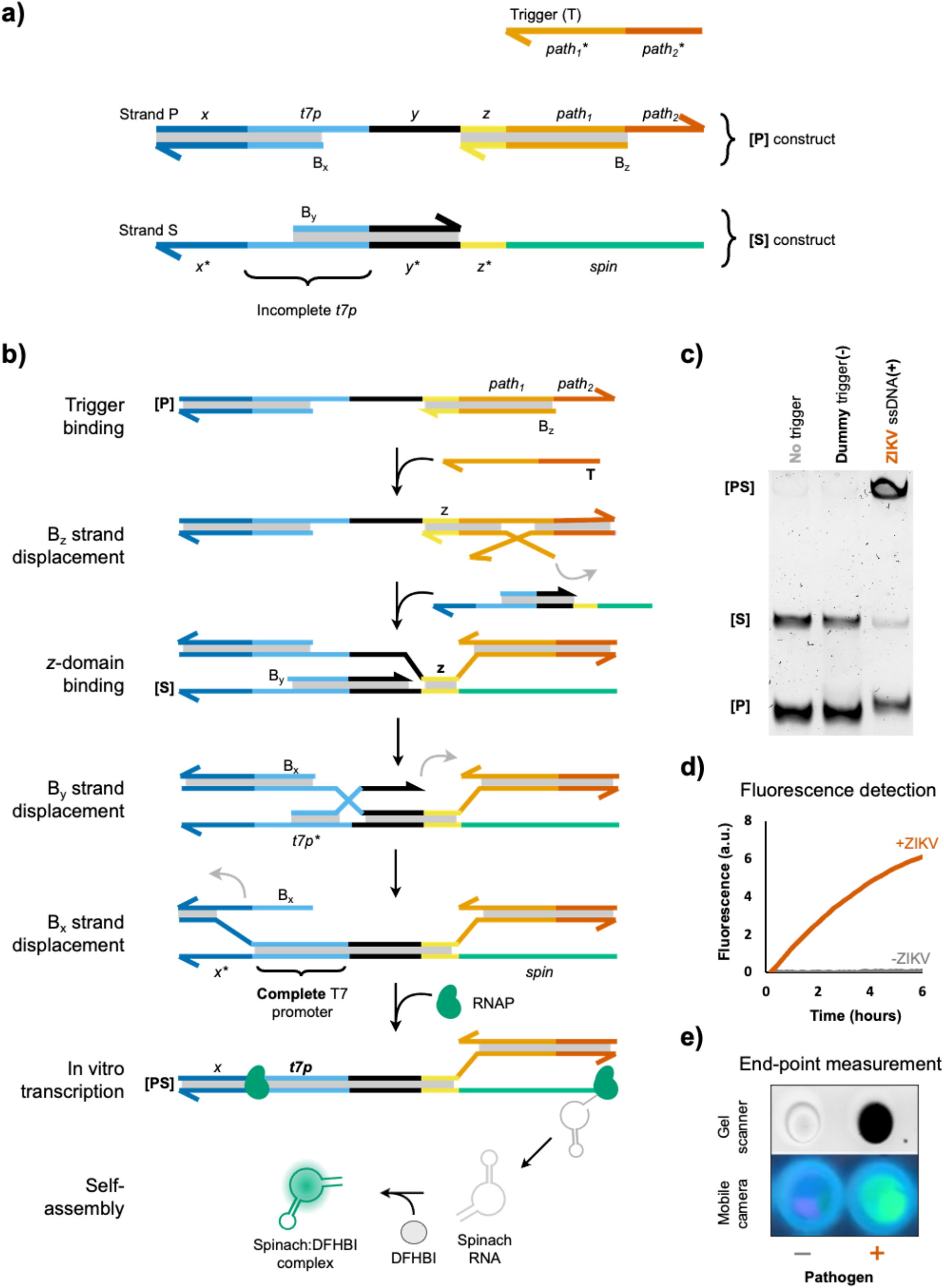
A nucleic acid target binds and thereby activates the thermodynamically spring-loaded gene switch, triggering a chain reaction that causes the expression of a fluorescent aptamer. Names of DNA constructs are bold and enclosed by square brackets (**[P], [S], [PS]**); individual DNA strands are denoted by capital letters (P, S, B_x_, etc.); sequence domains are written in lower case italics (*x, t7p, path*_*1*_, etc.). a) Design of **[P]** and **[S]** constructs. **[S]** contains a major strand, S, with a T7 promoter (*t7p*) domain upstream of a Spinach-encoding domain (*spin*). t7p is partially single stranded and hence cannot recruit T7 RNA polymerase (RNAP). **[P]** contains a major strand, P, that is largely complementary to S, but instead of spin it contains a pathogen-recognizing domain (*path*), which is partially double stranded (*path*_*1*_) and partially single stranded (*path*_*2*_). *path*_*2*_ serves as a toehold for the initial binding of a pathogen-specific trigger (T). The presence of blocking strands B_x_, B_y_, and B_z_ generates a large kinetic barrier for the initiation of the displacement cascade, thus preventing the formation of an active T7 promoter domain in the absence of a trigger strand. The sequence domains x and y are necessary to stabilize the attachment of B_x_ and B_y_, respectively, whilst domain z serves as the initial binding site between P and S strands. b) Designed mechanism for the triggered TMSD cascade, Spinach transcription, and self-assembly with the conditional fluorophore DFHBI. c) Typical PAGE gel showing that pathogen-specific sequence (ssDNA with ZIKV-specific sequence, 50 nM) triggers the strand displacement cascade. Small quantities of leakage product are detected in lanes 1 and 2. d) Time-dependent fluorescence in the presence and absence of the pathogen sequence. e) The presence or absence of a pathogen trigger can be detected on a laboratory gel scanner or a mobile phone camera.

In presence of a pathogen-specific nucleic acid sequence, a cascade of TMSD reactions is set into motion that ultimately leads to activation of the t7p domain and expression of a fluorescent signal (Figure 2b). The cascade is initiated by binding of the target sequence to the toehold *path*_*2*_, which allows displacement of B_z_. This first step exposes the *z* domain on **[P]**, which can subsequently bind to **[S]**, thus enabling the stepwise displacement of the remaining blocking strands, B_y_ and B_x_. The end product **[PS]** contains a double stranded *t7p* domain. T7 polymerase now starts transcribing a large number of Spinach aptamers that each bind and thereby light-up the conditional fluorophore 3,5-Difluoro-4-hydroxybenzylidene imidazolidinone (DFHBI)^48,49^.

### The assay creates a fluorescence signal that is detectable by a smartphone camera

As a first validation of the assay’s function, we incubated **[P]** and **[S]** in equimolar ratio and tested the response of the system in the presence and absence of its intended ssDNA trigger strand (T) at 50 nM concentration (Figure 2c, Supplementary Figure S1). Adding the trigger resulted in a high-molecular weight band that is consistent with the strand displacement product **[PS]**. Little strand displacement product was detected when the trigger was replaced by a dummy DNA strand with a random base sequence. Notably, the leakage product is distinguishable on PAGE from the trigger-induced product, as the leakage product still contains the B_z_ strand (Supplementary Figure S2). We next carried out the same reaction in the presence of T7 RNA polymerase and DFHBI and recorded the green fluorescence signal that is indicative of the expected Spinach-DFHBI complex. In agreement with the sensor’s design, a strong fluorescence signal was generated in the presence of T but not in trigger-free control samples (Figure 2d, e). Fluorescence measurements were recorded on a real-time qPCR instrument in order to capture the time-dependent profile of signal increase (Figure 2d). Importantly, the readout can also be performed as an end-point measurement, using either a laboratory gel scanner or an inexpensive smartphone camera (Figure 2e). The latter approach highlights the practicality of the assay for point-of-care detection, utilizing widely accessible personal equipment. A key benefit of the assay is the low fluorescence background of DFHBI, coupled with strong signal intensity resulting from the cumulative expression of Spinach aptamers over time. The successful expression of the RNA aptamer was also validated via PAGE (Supplementary Figure S2).

### TMSD leakage can be greatly reduced but not fully eliminated

Despite the sensor’s sequence-specific response, it produced minor quantities of leakage products (Figure 3a), precluding detection of sub-nanomolar analyte concentrations. To examine the leakage profile, we incubated equimolar mixtures of **[P]** and **[S]** at high concentration (200 nM each) for up to 48 hours and quantified the leakage by gel electrophoresis (Figure 3b).

**Figure 3.**
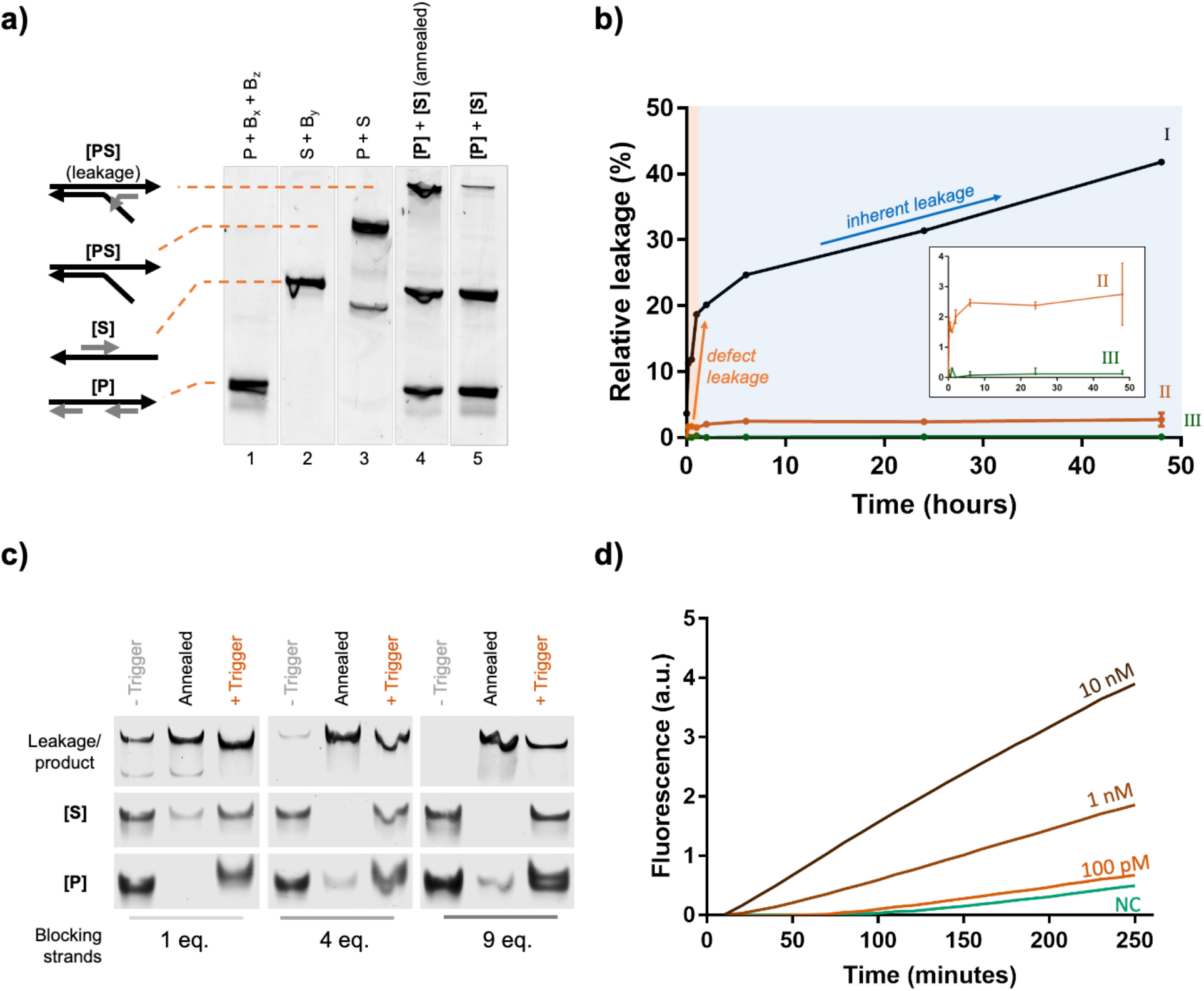
Reducing inherent and defect-related TMSD leakage is crucial for a sensitive sensor response. a) PAGE analysis of a typical leakage product after incubating **[P]** (lane 1) and **[S]** (lane 2) together for 30 minutes at room temperature (lane 5), and after quick annealing to 80°C (to simulate an excessive degree of leakage; lane 4), in comparison with the pure **[PS]** reference (lane 3). The leakage product is shifted with respect to the pure **[PS]** band, as B_z_ is still attached. b) Leakage profile of the TMSD cascade design versions 1, 2, and 3 (cf. Supplementary Note 2.2): (I) Design version 1 (with 200 nM construct concentration and 2 eq. blocking strands) reveals a fast initial leakage (defect-leakage), followed by a slow and continuous (inherent) leakage. (II) Design version 2 (with 20 nM construct concentration, 10 eq. blocking strands, and extended *x* and *y* domains) showed drastically reduced inherent and defect-related leakage rate. (III) In design version 3 (leakage-tolerant design; 20 nM construct concentration, 3 eq. blocking strands, biotinylated blocking strands), leakage was undetected after streptavidin bead capture. Error bars represent the standard deviation of 3 replicate experiments. Source gel images are shown in Supplementary Figures S3 and S4. c) PAGE analysis shows the reduction in leakage by introducing an increasing excess of blocking strands. All samples were incubated at 25 °C for 4 hours prior to PAGE analysis. d) Sensitivity assay with different concentrations of ssDNA oligonucleotide trigger revealed that the TMSD cascade (version 2) is able to detect trigger concentrations as low as 100 pM.

The data reveals two distinct leakage phases: an initial rapid increase that takes place within the first 60 minutes, consuming approximately 20% of **[P]** and **[S]** constructs; and a second phase that proceeds slowly but continuously for days (Figure 3b, trace I). An apparent stepwise leakage is typically associated with multiple distinct leakage pathways^32,34^. We hypothesized that physical defects in the system (e.g., ‘shortmer’ contaminations^52,53^ and incomplete construct assembly) are responsible for the fast initial phase, affecting approximately 20% of constructs (*defect leakage*). The second linear leakage phase suggested an inherent issue with the system’s design, likely permitting occasional binding of correctly assembled **[P]** to **[S]** constructs (enabled by fraying, breathing, and other thermal fluctuations) to spontaneously form **[PS]** (*inherent leakage*).

Aiming for a leakage-free system, we implemented several optimizations (Supplementary Note 2.2). These improvements deliberately excluded the use of additional oligonucleotide purification, which is costly and time-consuming^22,39,53^. Instead, we aimed for an inexpensive system that exhibits negligible inherent leakage while tolerating the existence of physical defects. The improvements included: (i) Extending the length of *x* and *y* domains from 10 to 15 nucleotides to increase the stability of blocking strand binding (Supplementary Figure S5). (ii) Adjusting the length of blocking strand overlaps (‘clamps’) to increase the tolerance for DNA fraying events^30,34^. (iii) Using a larger excess of blocking strands to allow for the in-situ replacement of blocking strand shortmers (Figure 3c). (iv) Reducing the constructs’ concentration from 200 nM to 20 nM to avoid unnecessarily frequent interactions between non-activated **[P]** and **[S]** (Supplementary Figure S6). Combined, these four improvements reduced the rate of inherent leakage by approximately 97% and defect-related leakage by approximately 92% (Figure 3b, trace II; Supplementary Table S5. Over the time course of 48 hours, over 97% of **[P]** and **[S]** constructs were not consumed by leakage processes.

### SARS-CoV-2 and ZIKV-derived nucleic acid sequences can be detected

To evaluate the performance and versatility of the improved assay, we exposed the TMSD reaction mixture to varying concentrations of pathogen-derived nucleic acid sequences and recorded the response of the Spinach aptamer fluorescence signal that was in-situ expressed by RNAP. We first tested a Zika-Virus-(ZIKV)-specific sequence and subsequently re-programmed the sensor to detect severe acute respiratory syndrome coronavirus 2 (SARS-CoV-2). This was achieved by replacing the ZIKV-specific *path* domain with a sequence that recognizes SARS-CoV-2’s N gene (Figure 4a). We tested the sensitivity for 33-nt long ssDNA targets, 33-nt ssRNA targets, and a 1.2 kb full-length mimic of the SARS-CoV-2 N-gene RNA. Raw fluorescence values were always normalized by the fluorescence of trigger-free reference sample. The critical limit (L_c_) for detecting ZIKV-derived DNA and RNA was 25 pM (500 amol, ∼5 pg) and 90 pM (1.8 fmol, ∼20 pg). Similar values were obtained for SARS-CoV-2 DNA and RNA, exhibiting L_c_ values of 75 pM (1.5 fmol, ∼15 pg) and 90 pM (1.8 fmol, ∼20 pg), respectively. No significant fluorescence signal increase was observed for negative control (NC) tests samples containing 800 ng of a cytoplasmic RNA reference mixture, demonstrating that the sensor is not triggered by a complex mixture of non-target nucleic acids (such as messenger- or ribosomal RNA) that is found in human cells. The detection of the 1.2 kb SARS-CoV-2 RNA was notably less sensitive than for the RNA oligonucleotides, exhibiting an L_c_ value of ∼1.4 nM (Figure 5d). The reduced sensitivity is likely due to secondary structures that are expected to exist in longer target strands^27^, thus lowering the rate of binding to the *path* domain.

**Figure 4.**
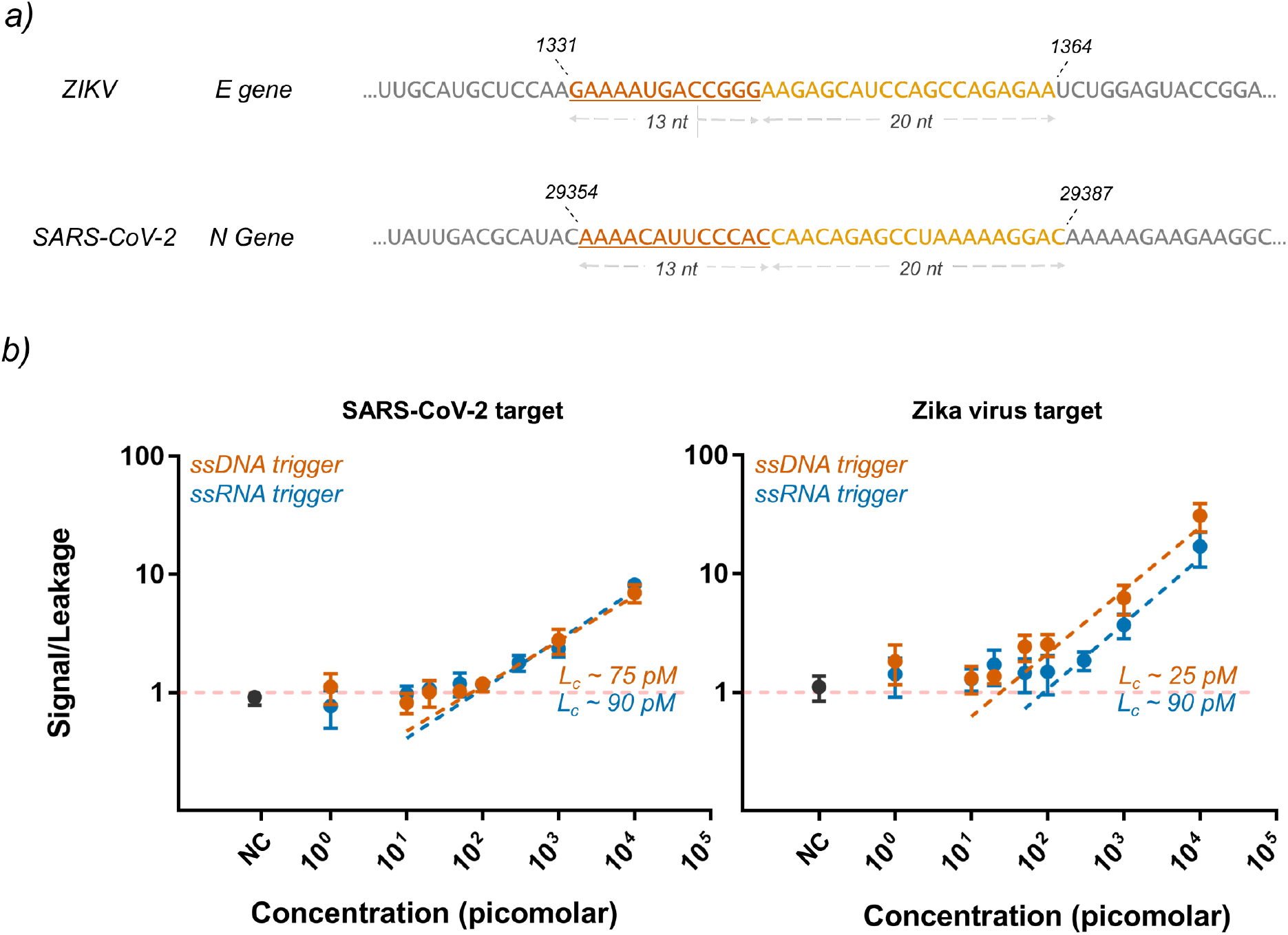
The reaction cascade responds to Zika Virus- and SARS-CoV-2-specific DNA and RNA sequences with similar sensitivity. a) Selected target regions for ZIKV and SARS-CoV-2 and their corresponding positions in the pathogen genome. The toehold-binding domain is underlined. b) Dependence of the normalized output signal intensity on the concentration of DNA and RNA targets specific to ZIKV and SARS-CoV-2. The L_c_ value was determined as the intersection of the linear regression with the threshold fluorescence increase, giving a quantitative measure for the theoretical detection limit of the assay. NC = negative control containing human cytoplasmic RNA. Individual plots with data analysis are shown in Supplementary Figure S8. The data was collected from three independent experiments for each target type.

**Figure 5.**
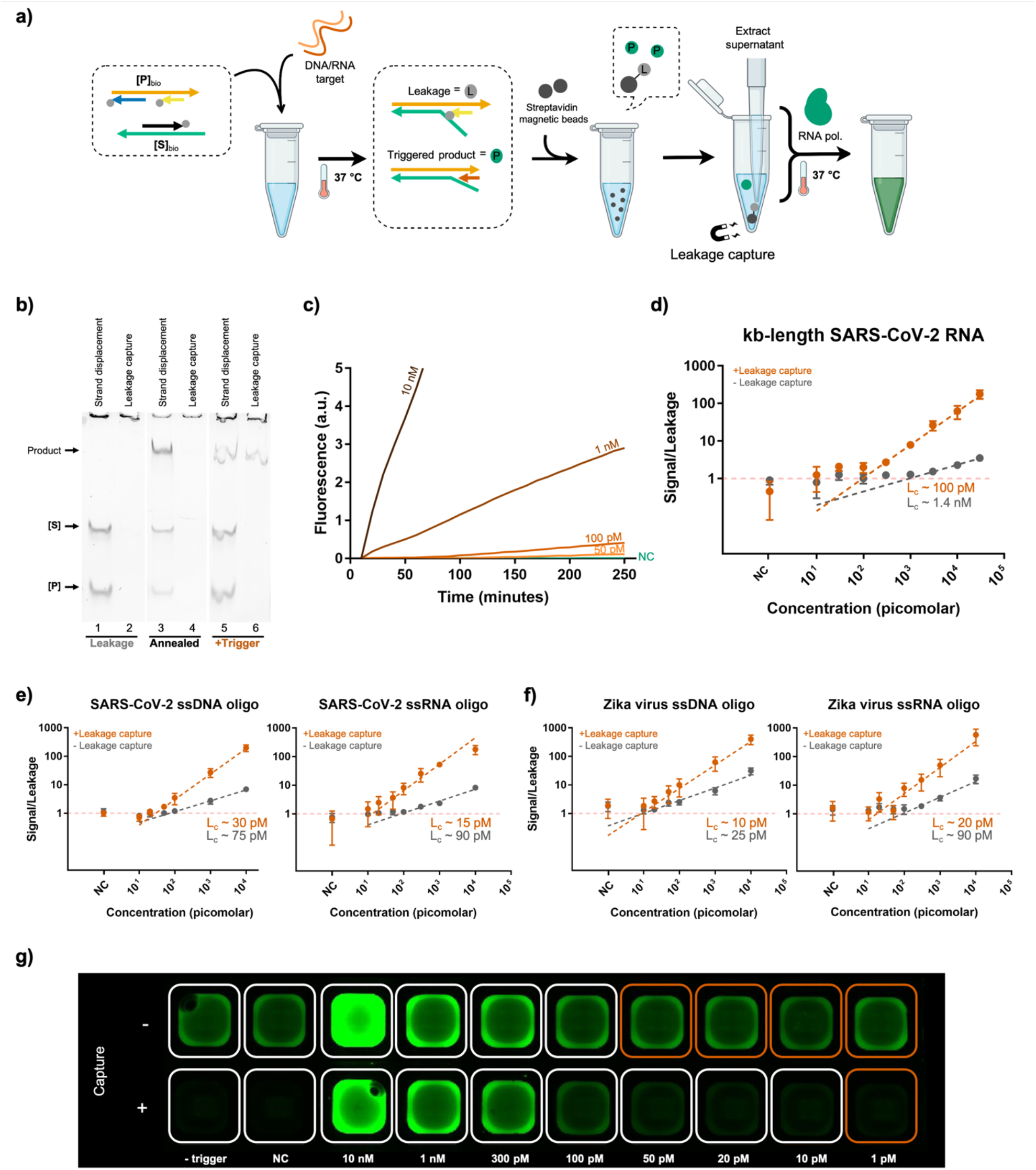
Leakage-tolerance is achieved through the selective capture of cascade components and byproducts prior to Spinach expression. a) Schematic representation of the assay with leakage capture (cascade version 3). Streptavidin beads selectively pull down **[P]**_bio_, **[S]**_bio_, and the leakage product, but not the correctly activated product **[PS]**. b) PAGE analysis shows the cascade components after a 30-minute incubation period, before vs. after leakage capture. Lanes 1, 2: samples incubated at 37°C in the absence of trigger strand. Lanes 3, 4: samples briefly annealed to 80°C in the absence of trigger strand to simulate excessive leakage. Lanes 5, 6: samples incubated at 37°C in the presence of trigger strand. c) Representative plots of raw fluorescence measurements, demonstrating the detection at different trigger concentrations. The unimpeded growth of the fluorescence signal indicates continuous expression of Spinach aptamers. d)-f) Plots of the signal-to-leakage ratio vs. trigger concentration for the leakage-reduced assay (cascade version 2) and the leakage-tolerant assay (cascade version 3). Leakage-capture improves the detection of 1.2 kb long SARS-CoV-2 RNA (d), 33-nt DNA and RNA oligonucleotides derived from SARS-CoV-2 (e), and Zika virus (f). The data for panels d, e, and f was collected from three replicate experiments for each target type. Individual plots with data analysis are shown in Supplementary Figure S9 and S10. g) End-point fluorescence image of a SARS-CoV-2 assay plate demonstrates that leakage capture reduces the background signal and thus lowers the concentration of detectable target strands. The wells marked in red contain test samples with background-level fluorescence intensity.

### Selective capture of TMSD reactants and byproducts makes the assay leakage-tolerant and more sensitive

Even though the optimizations to the sensor had demonstrated initial sensitivity improvements (Figure 3 & 4), exhaustive attempts to further reduce leakage through TMSD design optimizations were unsuccessful. We reasoned that the large diversity of leakage pathways—some involving shortmers, branched oligos, or internal deletion products—made further optimizations challenging. Instead of seeking perfect leakage suppression, we therefore aimed at making the system *leakage-tolerant* through selective capture of any leakage product that is emerging prior to signal amplification. Independent of the exact leakage pathway, the final leakage product was expected to be **[PS]** with B_z_ still bound to the *path* domain (Figure 5a and Figure 3a). To allow selective capture, we replaced all blocking strands with their 3’-biotinylated analogs. This modification allowed quantitative pulldown of the leakage product with Streptavidin-coated magnetic beads (Figure 5a). The bead pulldown also removes the biotinylated constructs (**[P]**_bio_ and **[S]**_bio_), thus precluding the formation of new leakage products during subsequent sample incubation.

For experimental validation, we compared the products of 30-minute **[P]**_bio_ + **[S]**_bio_ TMSD reactions before and after streptavidin bead pulldown. The TMSD cascade was carried out with and without ssDNA trigger oligos, and along with a thermally annealed reaction that simulated an excessive degree of leakage. PAGE analysis confirmed that Streptavidin beads captured and removed **[P]**_bio_, **[S]**_bio_, and leakage product with 96-98% efficiency, while leaving only the trigger-activated **[PS]** construct in the supernatant (Figure 5b, Supplementary Table S6). The supernatant showed greatly reduced expression of Spinach aptamer, which was undetectable by PAGE.

Fluorescence measurements confirmed the greatly reduced background signal of the leakage-tolerant assay (Figure 3b, trace III), as compared to the merely leakage-reduced assay (Figure 3b, trace II). In the presence of a trigger, we observed near-ideal linearly growing amplification, suggesting that once activated, Spinach expression can proceed unimpeded for hours (Figure 5c). The leakage-tolerant assay showed consistently high sensitivity (L_c_: 10-30 pM) for ssDNA and ssRNA oligos of both ZIKV and SARS-CoV-2, outperforming the leakage-reduced assay for all target types (Figure 5e, f). We also observed a remarkable enhancement in performance for detecting the 1.2 kB SARS-CoV-2 RNA mimic, where the L_c_ was improved from 1.4 nM to 100 pM (Figure 5d). While the detection sensitivity was assessed based on time-resolved fluorescence measurements, simple end-point measurements were found to be similarly suitable. The fluorescence images further demonstrate the increased assay sensitivity due to a much reduced background signal in the leakage-tolerant mechanism (Figure 5g). Overall, the results validate the benefit of a cascade design that allows for selective leakage capture to achieve a leakage-tolerant detection workflow with increased sensitivity.

## DISCUSSION

In summary, this work presents a programmable and isothermal nanosensor that is triggered by single-stranded nucleic acids. The underlying TMSD reaction cascade serves as a molecular switch controlling the in-situ expression of a fluorescent aptamer. Due to its modularity, the cascade can be quickly adapted to detect arbitrary targets. Both DNA and RNA can be detected at picomolar concentrations without requiring target amplification, reverse transcription, or any other enzymatic pretreatment. We validated the sensor with sequences derived from Zika virus and SARS-CoV-2 genomes (Figure 4b, Figure 5e, f), demonstrating that its performance can be seamlessly transferred to a different target without the need for experimental sequence screening.

The sensor’s architecture is functionally similar to previously reported genelet designs^33,35,36,54^, but exhibits two key differences: firstly, once activated, the *t7p* domain is entirely double stranded and does not contain any nicks (Figure 2b). The intactness of the promotor enables more efficient transcription and prevents defective RNA displacement, which can cause spurious inactivation of the transcription template in genelets^51^. The cascade design therefore enables continuous and nearly unimpeded aptamer expression from **[PS]** over many hours, leading to a strong output signal (Figure 5c). Secondly, while the input for the canonical genelet design contains a sequence fragment of the t7 promoter, the TMSD cascade presented herein does not impose any constraint on the input sequence. Therefore, the sensor can be easily re-programmed to accept any single-stranded nucleic acid sequence as an input and produce an arbitrary RNA sequence as an amplified output.

Initial designs of our sensor (versions 0 and 1) suffered from notable strand displacement leakage, an obstacle that is frequently observed in TMSD cascades^26,28–34^. Our investigation into the leakage dynamics guided the development of a strongly leakage-reduced (version 2) and ultimately leakage-tolerant (version 3) assay. The optimized sensor shows zero-background fluorescence, not only in the absence of any trigger but also when random DNA oligonucleotides or complex human cytoplasmic RNA mixtures are present in the sample.

Due to its strong fluorescence and low background signal, the sensitivity of the leakage-tolerant design is relatively high. Its critical concentration for response onset lies between 10 to 100 pM (in 10–20 μL reaction volume), with a minimal detectable total RNA amount of as little as 200 attomole (∼10^8^ copies). By comparison, clinical samples taken from the upper respiratory tract of COVID-19-positive patients contain 10^7^ to 10^11^ viral genome copies per milliliter^55^. For ZIKV-positive patients, 10^8^ to 10^9^ viral genome copies can be found in semen (but only as little as 10^3^ copies per milliliter in blood or urine)^56^. Importantly, the isothermally generated fluorescence output can be measured at the endpoint, by using either a plate reader or a mobile phone camera (Figure 2d, e), making it applicable at the point of care. Despite its encouraging performance characteristics and advantages for POC detection, the cascade’s sensitivity is not yet on par with well-established enzymatic amplification reactions like PCR or LAMP, for which detection of just a few genome copies is achievable^1^. There are several possible ways to further lower the system’s detection threshold: firstly, the sensitivity of cascade version 3 is not limited by TMSD leakage but rather by the total amount of fluorescent signal generated. Thus, we expect that higher sensitivities can be achieved by increasing the output expression rate (e.g., with a high-performance T7 promoter^57^) or by expressing a brighter aptamer (e.g., Broccoli^58^). Secondly, before activating the TMSD cascade, target sequences could be enriched from clinical samples prior to the TMSD cascade by employing recently developed pulldown polymers^59,60^. Thirdly, as the current system exhibits linear growth kinetics, an additional downstream reaction could be added to provide more rapid quadratic or even exponential signal growth^16^.

Future developments will widen the system’s scope in diagnostic applications, for instance for the detection of small non-coding RNA, or the discrimination of single-nucleotide variants^21^. Beyond diagnostics, we expect that this gene switch will serve as a new primitive for high-fidelity TMSD reaction networks, DNA-based signal amplification, and leakage-tolerant molecular computations^61–65^. While this TMSD cascade has been validated in solution, we envision that it can be assembled onto DNA-functionalized polymer scaffolds^17^ to endow bio-interfacing materials (e.g., dynamic cell culture matrices^66^) with sensory functions.

## ACKNOWLEDGEMENTS

We thank Prof. Dr. Sun Hur and Prof. Dr. Xin Mu (Harvard Medical School) for providing cytoplasmic RNA. We thank Prof. Dr. Alexander Dalpke, Rayan Suliman, and Leonie Wilke (Universitätsklinikum Carl Gustav Carus Dresden) for providing a SARS-CoV-2 RNA reference and Dr. Beatrise Berzina for assistance with an RNA quantification experiment. We further thank Dr. Michele Marass and Dr. Agnes Kriegne Toth-Petroczy for giving valuable feedback on the paper draft. Parts of Figure 1 and Figure 5a were created with BioRender.com.

