## Supplementary Information for "A spring-loaded and leakage-tolerant synthetic gene switch for in-vitro detection of DNA and RNA"

**1 SUPPLEMENTARY METHODS 2**

1.1 INITIAL VALIDATION OF TMSD CASCADE FUNCTION AND IDENTIFICATION OF LEAKAGE 2
1.2 INITIAL VALIDATION OF TRIGGER-SPECIFIC RESPONSE USING STRAND DISPLACEMENT ASSAY 2
1.3 INITIAL VALIDATION USING FLUORESCENCE ASSAY 2
1.4 TRACING THE KINETICS OF LEAKAGE IN DIFFERENT TMSD CASCADES 3
1.5 DENATURING POLYACRYLAMIDE GEL ELECTROPHORESIS (PAGE) ANALYSIS 3
1.6 EXPLORING LEAKAGE REDUCTION WITH DIFFERENT EXCESSES OF BLOCKING STRANDS 4
1.7 VALIDATING LEAKAGE-CAPTURE USING STRAND DISPLACEMENT ASSAY 4

**2 SUPPLEMENTARY NOTES 5**

*2.1* PRELIMINARY DESIGN OF THE TMSD CASCADE 5
*2.2* REFINED DESIGNS OF THE TMSD CASCADE 6
2.3 WORKFLOW FOR IDENTIFYING A SUITABLE TARGET SEQUENCE IN A PATHOGEN GENOME 7

**3 SUPPLEMENTARY FIGURES 10**

**4 SUPPLEMENTARY TABLES 17**

**5 SUPPLEMENTARY REFERENCES 19**

### 1 Supplementary Methods

#### 1.1 Initial validation of TMSD cascade function and identification of leakage

The initial validation of strand displacement and preliminary identification of leakage were conducted using **[P]** and **[S]** constructs from preliminary designs. Strands from giving TMSD cascade 0 were employed to assemble the constructs in the annealing buffer. They were assembled at 500 nM concentrations with 3 equivalents(eq.) blocking strands using the thermal annealing protocol. The reaction was conducted relative to 50 nM construct concentration. Different DNA components were mixed in a reaction mix composed of reaction buffer from HiScribe T7 High Yield RNA Synthesis Kit (New England Biolabs, Cat. # E2040), supplemented with 200 mM KCl. The samples were incubated at 37 °C for 10 minutes, and subsequently analyzed on a 10% native PAGE gel.

#### 1.2 Initial validation of trigger-specific response using strand displacement assay

The experiment with trigger-initiated strand displacement was conducted with TMSD Cascade 1_10nt. The constructs were assembled in an annealing buffer at 200 nM concentrations with 3 eq. blocking strands using the thermal annealing protocol. The strand displacement reaction was conducted with the constructs at 20 nM concentration. 33-nt long ZIKV_ssDNA and 60-nt long Dummy_ssDNA were used as ZIKV and dummy triggers respectively at 50 nM concentration. Different components were assembled in a reaction mix composed of 1X reaction buffer from HiScribe T7 High Yield RNA Synthesis Kit (New England Biolabs, Cat. # E2040), supplemented with 200 mM KCl. The samples were incubated at 37 °C for 10 minutes and subsequently analyzed on an 8% native PAGE gel.

#### 1.3 Initial validation using fluorescence assay

The initial fluorescence assays were conducted using the HiScribe T7 High Yield RNA Synthesis Kit (NEB, Cat. # E2040S) at a 20 µl reaction scale using sequences from the previous section. The reaction master mix was prepared by mixing the following volumes of components per sample: 1.5 µl 10X Reaction buffer, 2 µl KCl (2 M), 1 µl DFHBI (400 µM in DMSO), 3.5 µl Nuclease-free water, and 6 µl rNTPs (100 mM each). Then the required DNA constructs (20 nM final), trigger (50 nM), and nuclease-free water were added to a final volume of 18 µl, and incubated at 37 °C for 30 mins. Then, each sample was split into two tubes of 9 µl each, where in one set, 1 µl T7 RNA polymerase was added, and in the other just the enzyme storage buffer (100 mM NaCl, 50 mM Tris-HCl, 1 mM EDTA, 50% glycerol). All the samples were transferred on a white qPCR plate (Sarstedt, #72.1981.232) and run on a real-time qPCR instrument (Biorad, CFX96) for over 4 hours in 10-minute cycles. Each cycle involved a 9-minute incubation at 37 °C, followed by a 1-minute plate read at 20 °C using FAM mode. After the experiment, the samples were run on a 10% native PAGE gel.

For end-point measurements, one sample with trigger, and one sample without trigger were transferred on transparent qPCR plates and observed using Typhoon 9500 (GE Healthcare Life Sciences) using a blue LD laser (excitation at 473 nm), BPB filter, 400 Volts setting, and with the resolution of 50 µm/pixel. Alternatively, the same plates were visualized on the mobile phone camera of One Plus 6 after illuminating the plates with a portable general-purpose laboratory UV lamp (8W).

#### 1.4 Tracing the kinetics of leakage in different TMSD cascades

To study the leakage kinetics, the reaction master mix (8 µl per sample) was prepared without **[P]** and **[S]** constructs, and aliquoted to the required volume in pre-marked reaction tubes while accounting for the addition of the constructs later (1 µl each). The experimental time points were planned by backtracking the sample addition time using PAGE analysis as the reference. For 48-hour experiments, first, the 48-hour sample was prepared by adding the constructs in the pre-marked and pre-filled reaction tubes and transferring them to a PCR instrument set at the required temperature. Subsequently, other reactions were started at stipulated time points by adding the constructs. At time t=0 hours, the samples were run on a 6% native PAGE gel.

**Trace I** was studied with sequences from TMSD Cascade 1_10nt. The leakage was studied at construct concentrations of 200 nM, and 2 eq. blocking. The incubation was conducted at 37 °C, n=1. The percentage of leakage at different time points was quantified using the sum of the intensity of all bands in a lane as a reference using the equation:

$$leakage\ (\%) = \frac{intensity_{[\boldsymbol{PS}]}}{intensity_{[lane]}}\ x100$$

**Trace II** employed sequences from TMSD Cascade 2_15nt_ZIKV, studying the leakage at 20 nM construct concentration, 10 eq. blocking strand and 25 °C reaction temperature, n=3.

**Trace III** used sequences P3, S3, and biotin-TEG tagged blocking strands: Bx3(bio), By3(bio), and Bz3(bio), from TMSD Cascade 2_15nt_ZIKV. After incubating the constructs at 20 nM construct concentration, 3 eq. blocking strand and 37 °C reaction temperature. For leakage capture, the resulting displacement product was added to pre-washed streptavidin-coated magnetic beads Dynabeads™ MyOne™ Streptavidin T1 (Thermo Scientific, #65601). The samples were incubated with magnetic beads for 30 minutes with a rotator-vortexer (neolab, Cat. #7-0045) using a custom 3D printer shaker attachment and then captured using a custom 3D printed magnetic separation stand. The supernatant was extracted, n=3.

For **Trace II** and **Trace III,** an additional annealed sample was prepared by thermally annealing both constructs in the reaction buffer as a quantitation reference, representing ~100% leakage. This reference was chosen because the efficiency of the annealed product was quantitative. The percentage of leakage was calculated as:

$$leakage\ (\%) = \frac{intensity_{[\boldsymbol{PS}]}}{intensity_{[annealed]}}\ x100$$

#### 1.5 Denaturing polyacrylamide gel electrophoresis (PAGE) analysis

A 4% denaturing polyacrylamide gel was cast using the SequaGel-UreaGel System (National Diagnostics, cat. #EC833). 9 µl of formamide loading buffer (98% Formamide(aq.), 5 mM EDTA, 1.7 mM Tris base, 0.02 wt % BPB, pH 8.0) was added to 1 µl RNA sample and heat treated (80 °C for 2 minutes, 4 °C for 5 minutes). The gels were loaded and run at 120 V using Consort EV265 power supply, stained using 1X

SYBR™ Gold, and imaged on Typhoon FLA 9500 (GE Healthcare Life Sciences) using a blue LD laser (excitation at 473 nm), BPB filter, 400 Volts setting, and with the resolution of 50 µm/pixel.

#### 1.6 Exploring leakage reduction with different excesses of blocking strands

To assemble the constructs, strands from TMSD Cascade 2_15nt_COVID were used. Different constructs were assembled using 2, 5, and 10 eq. the concentration of blocking strands. The constructs were assembled at 100 nM concentration in an annealing mix (40 mM Tris-HCl, pH 7.9, 12 mM $MgCl_2$, 150 mM KCl) using the annealing protocol. 20 nM COVID_ssDNA was used as the trigger. The respective constructs were added to the reaction mix (40 mM Tris-HCl, pH 7.9, 12 mM $MgCl_2$, 150 mM KCl) at 20 µl reaction volume and incubated at 25 °C for 4 hours. Meanwhile, the tubes belonging to annealed samples were re-annealed with both constructs using the annealing protocol. The samples were run on an 8% native PAGE gel.

#### 1.7 Validating leakage-capture using strand displacement assay

Constructs were assembled at 200 nM concentration in an annealing buffer using sequences from TMSD Cascade 2_15nt_COVID, 3 eq. concentration of biotin-TEG capped blocking strands. The constructs and COVID_ssDNA trigger were added to the reaction buffer (40 mM Tris-HCl, pH 7.9, 12 mM $MgCl_2$, 150 mM KCl) and incubated at 37 °C for 30 minutes. In the meanwhile, the tubes belonging to annealed samples were thermally re-annealed by using annealing protocol. After incubation, the samples were added to pre-washed streptavidin-coated magnetic beads, and incubated at room temperature with shaking for 30 minutes. After the capture of magnetic beads, the supernatant from all the samples were run on an 8% native PAGE gel.

### 2 Supplementary Notes

#### *2.1* Preliminary design of the TMSD cascade

**Cascade version 0.** Constructs **[P]** and **[S]** were initially designed as shown in the scheme below. The *x*-, *y*-, and *z*- domains were 10, 10, and 8 nucleotides long, respectively. In this concept, the pathogen-target-binding toehold (*$path_2$*) was designed to lie on blocking strand $B_z$. This design, termed "Version 0", was used for preliminary internal testing.

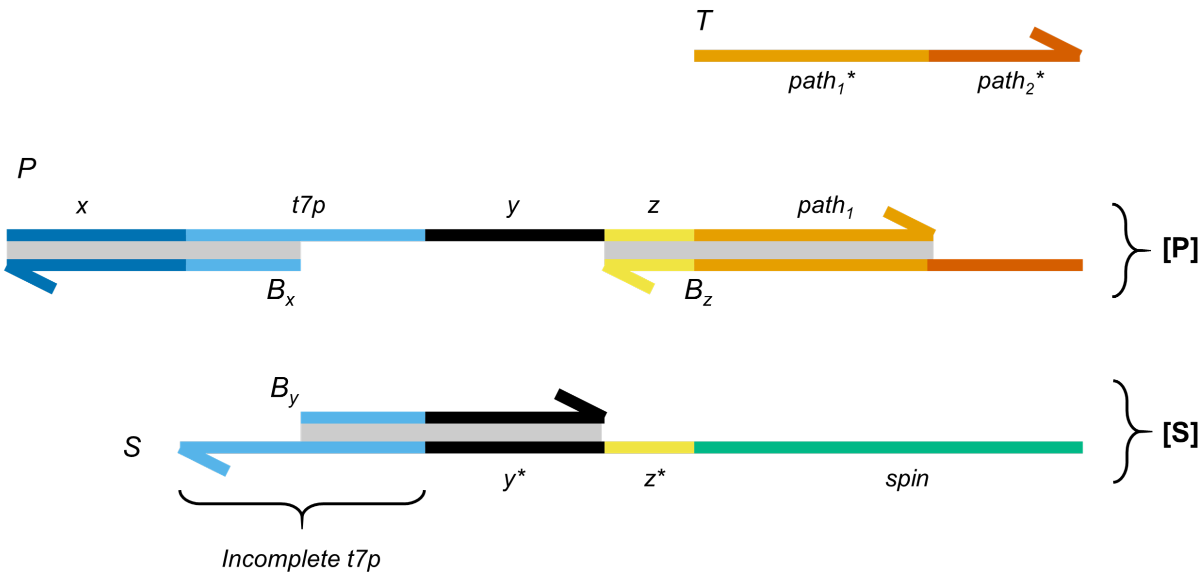


The NUPACK design tool was used to generate specific sequences for the individual domains, using the following script in DU+ notation. In this design, the 30-nt *path* domain was generated by extending the RT-qPCR forward primer sequence (5'-AATGGGAAGGAAAGAAGAGG-3') used in Lamb et al.[1]. We chose the *$path_1$* domain to be 20 nt long, which yielded a 10-nt *$path_2$* domain as the toehold. Salt concentrations were set to 125 mM NaCl and 5 mM $MgCl_2$. The choice of sequences from a list of candidates was based on minimal undesired secondary structures, each nucleobase had to be represented to at least 15% in the sequence, and the GC content had to be within the range of 40-60%.

**Code 1.** Initial NUPACK design for the TMSD system.

```
material = dna
temperature[C] = 37.0 # optional units: C (default) or K
trials = 10
sodium= 0.125
magnesium = 0.005
#
# target structures
#syntax
#structure signal1 = U36 <write the bindings DU+ notation>
structure PS= U10 D38 (U20+U98)
structure PBX=D20 U48
structure PBY=U20 D20 U28
structure SBX= U136 + U20
structure SBY= U136+ U20
structure strucBX= U20
structure strucBY= U20
structure strucP= U68
```

```
structure strucS= U136
#
# sequence domains
#syntax
#domain <domain_name>= N30 <sequence names>
domain X= N10
domain Y= N10
domain Z= N8
domain T71= TAATACGACT
domain T72= CACTATAGGG
domain Path= AATGGGAAGGAAAGAAGAGG

domain
SPIN= GACGCGACTAGTTACGGAGCTCACACTCTACTCAACAAGCTGCACTGCCGAAGCAGCCACACCTGGACCCGTCCTTCACCATTTCATTC
AGTTGCGTC

# strands (optional, used for threading sequence information and for displaying results)
# syntax:
# strand <strand_name>= domain1 domain1* domain2
strand P= X T71 T72 Y Z Path
strand S= SPIN Z* Y* T72* T71*
strand BX= T71* X*
strand BY= Y* T72*
#
# thread strand sequence information onto target structures
# syntax
#<structurename>.seq= <strand_name>
PS.seq= P S
PCX.seq= P BX
PCY.seq= P BY
SBX.seq= S BX
SBY.seq= S BY
strucBX.seq= BX
strucBY.seq= BY
strucP.seq= P
strucS.seq= S
#
# prevent sequence patterns
#
prevent = AAAA, CCCC, GGGG, UUUU, KKKKKK, MMMMMM, RRRRRR, SSSSSS, WWWWWW, YYYYYY
```

#### *2.2* Refined designs of the TMSD cascade

**Cascade version 1.** Preliminary experiments with cascade version 0 indicated that it is advantageous to include $path_2$ on strand P, rather than $B_z$, since any excess of free blocking strands sequester target nucleic acids and thus reduces the assay's sensitivity. Furthermore, to increase the thermodynamic stability of the activated construct **[PS]**, the S strand was extended to include the x-domain. We also appended six extra thymine bases to the 5' end of both P and S strands, to eliminate the possibility that difficult-to-remove shortmer contaminations[2] (n-1, n-2, … n-6) would lack parts of their functional domains. Lastly, the toehold domain was slightly extended from 10 to 13 nt in length to increase the stability of the initial pathogen-sequence-bound complex.

**Cascade version 2.** Further optimizations were implemented to drastically reduce leakage, yielding version 2 of the assay. To increase the stability of blocking strand binding, the *x*- and *y*- domains were extended to 15 nt (Supplementary Figure S5). To reduce leakage that is a result of fraying[3,4], additional clamping domains were introduced (Supplementary Figure S6). Moreover, the excess of blocking strands was increased from 2-3 to 10 equivalents. The larger excess of blocking strands was intended to facilitate the efficient in-situ replacement of shortmer blocking strands (i.e. oligonucleotide synthesis byproducts) by full-length blocking strands. Furthermore, we reduced the final concentration of **[P]** and **[S]** in the detection solution from 200 nM to 20 nM, to avoid unnecessarily frequent interactions between constructs.

**Cascade version 3.** To make the system leakage tolerant, we added 3'-Biotin modifications to all blocking strands. This upgrade enabled the selective pulldown of all reaction components and leakage byproducts via streptavidin bead pulldown. The leakage-tolerant mechanism also allowed us to reduce the concentration of blocking strands from 10 to 3 eq., which was expected to favorably affect the cascade's reaction rate and efficiency. All other parameters of the TMSD cascade are identical to cascade version 2.

The stepwise improvements from cascade 1 to 3 are reflected in the main text Table 1, Supplementary Table S2, and Supplementary Figure S6. The effect on leakage is experimentally illustrated in the main text Figure 3b.

#### 2.3 Workflow for identifying a suitable target sequence in a pathogen genome

The objective was to develop a workflow to identify an ideal 33-nt region from a pathogen genome to be targeted by a TMSD-based detection cascade. As no thermal or chemical denaturation takes place in our assay, the target region should be mostly single-stranded at 37°C to trigger the sensor. Our approach involved dividing the target RNA sequence into fragments, and identifying parts that are predicted to have minimal secondary structure. Subsequently, candidate sequences were screened against undesired non-specific interactions with both the **[S]** and the **[P]** construct.

Firstly, the N gene reference sequence was used as identified from the Wuhan-Hu-1 isolate (GenBank: MN908947.3) and the 1260 nts long sequence was sliced into 200 nt fragments with 20 nt overlaps, yielding 7 fragments. The overlaps were used to consider the influence of immediate adjoining regions to form secondary structures. We conducted NUPACK simulations at 37 °C to predict the possible structure of RNA sequence and identify linear regions (Figure SN1a). Subsequently, the linear regions (Figure SN1b) were chosen manually following the dot parentheses notation[5]. These fragments underwent a more stringent screening process for secondary structures, which included analyzing the sequences for pseudoknots (Figure SN1c).

a)

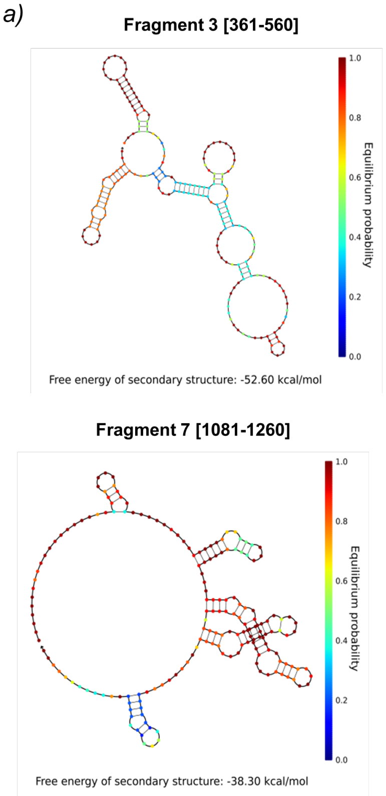


b)

| # | Fragment | Start | End | Size [nt] | Sequence |
|---|---|---|---|---|---|
| 1 | 1 | 1 | 25 | 25 | AUGUCUGAUAAUGGACCCCAAAAUC |
| 2 | 1 | 52 | 99 | 48 | GGUGGACCCUCAGAUUCAACUGGCAGUAACCAGAAUGGAGAACGCAGU |
| 3 | 1 | 129 | 176 | 48 | AGGUUUACCCAAUAAUACUGCGUCUUGGUUCACCGCUCUCACUCAACA |
| 4 | 2 | 181 | 220 | 40 | AAGGAAGACCUUAAAUUCCCUCGAGGACAAGGCGUUCCAA |
| 5 | 3 | 398 | 465 | 68 | UUGCAACUGAGGGAGCCUUGAAUACACCAAAAGAUCACAUUGGCACCCGCAAUCCUGCUAACAAUGCU |
| 6 | 3 | 476 | 514 | 39 | UACAACUUCCUCAAGGAACAACAUUGCCAAAAGGCUUCU |
| 7 | 4 | 541 | 576 | 36 | CAAGCCUCUUCUCGUUCCUCAUCACGUAGUCGCAAC |
| 8 | 5 | 721 | 752 | 32 | CAACAAGGCCAAACUGUCACUAAGAAAUCUGC |
| 9 | 5 | 894 | 920 | 27 | CAAACAUUGGCCGCAAAUUGCACAAUU |
| 10 | 6 | 1175 | 1100 | 26 | GCAUACAAAACAUUCCCACCAACAGA |
| 11 | 7 | 1081 | 1122 | 42 | AAAACAUUCCCACCAACAGAGCCUAAAAAGGACAAAAAGAAG |
| 12 | 7 | 1123 | 1260 | 38 | AACAAUCCAUGAGCAGUGCUGACUCAACUCAGGCCUAA |

c)

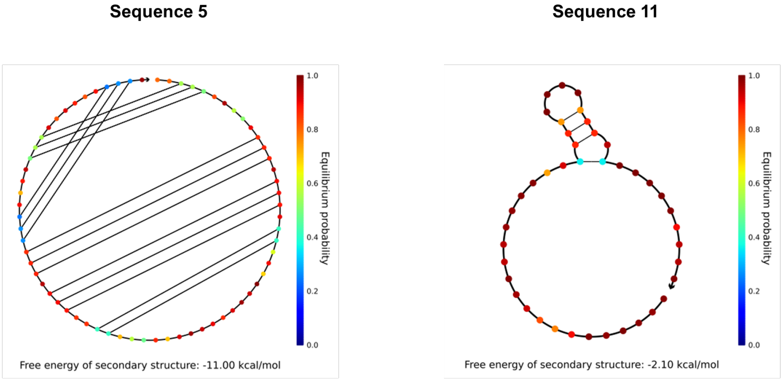


**Figure SN1.** Preliminary screening of suitable pathogen recognition site using NUPACK analysis a) NUPACK simulations showing analysis of 2 *N* gene fragments to identify regions with possible linear sites. b) Candidates for pathogen recognition sites after fragment analysis c) Analysis of 2 example SARS-CoV-2 target candidate sequences that were screened against secondary structures (including pseudoknots).

The selection criteria mandated that the sequence length exceeds 33 nucleotides, exhibit minimal secondary structures, and allow no more than three consecutive base pairs. Subsequently, the candidates were also screened against interactions with the rest of the SARS-CoV-2 genome. Out of 12 initial candidate sequences, only two of them, #5 and #11, met all criteria. They were trimmed down into different combinations of 33 nt regions, screened for secondary structures and sequence complexity via GC content, which yielded two final hits (Figure SN2a).

*a)*

| # | Sequence | Length | Structure | Reverse complementary DNA |
|---|---|---|---|---|
| 1 | UUGCAACUGAGGGAGCCUUGAAUACACCAAAAG | 33 | 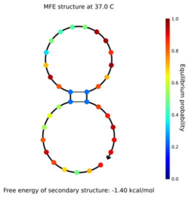  | CTTTTGGTGTATTCAAGGCTCCCTCAGTTGCAA |
| 2 | AAAACAUUCCCACCAACAGAGCCUAAAAAGGAC | 33 | 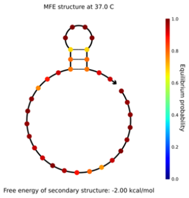  | GTCCTTTTTAGGCTCTGTTGGTGGGAATGTTTT |

*b)*

| | 1 EQ | | | 2 EQ | | | 5 EQ | | | 20 EQ | | |
|---|---|---|---|---|---|---|---|---|---|---|---|---|
| | 4 °C | 25 °C | 37 °C | 4 °C | 25 °C | 37 °C | 4 °C | 25 °C | 37 °C | 4 °C | 25 °C | 37 °C |
| 10 nM | 100 | 100 | 100 | 100 | 100 | 100 | 100 | 99 | 99 | | 97 | 97 |
| 20 nM | 100 | 100 | 100 | 100 | 100 | 100 | 100 | 95 | 95 | 100 | 95 | 95 |
| 200 nM | 100 | 100 | 100 | 100 | 100 | 100 | 100 | 100 | 95 | 100 | 95 | 90 |
| 500 nM | 100 | 100 | 100 | 100 | 98 | 98 | 100 | 98 | 96 | 98 | 92 | 88 |
| 1000 nM | 100 | 99 | 99 | 99 | 98 | 98 | 99 | 97 | 95 | 98 | 92 | 86 |

*c)*

| | 1 EQ | | 2 EQ | | 5 EQ | | 20 EQ | |
|---|---|---|---|---|---|---|---|---|
| | 25 °C | 37 °C | 25 °C | 37 °C | 25 °C | 37 °C | 25 °C | 37 °C |
| 10 nM | 100 | 100 | 100 | 100 | 100 | 100 | 100 | 100 |
| 20 nM | 100 | 100 | 100 | 100 | 100 | 100 | 100 | 100 |
| 50 nM | 100 | 100 | 100 | 100 | 100 | 100 | 100 | 100 |

**Figure SN2.** Integrating truncated linear regions with the universal template. a) Truncated linear regions showing their predicted structure, and reverse complementary region (DNA). The sequences predict high efficiency (%) of strand displacement simulated in the b) absence and c) presence of pathogen trigger. The simulations were done under experimental conditions of 50 mM $Na^{+}$, 36 mM $Mg^{2+}$, different equivalents of blocking strands, and at different temperatures to predict the system's equilibrium.

The respective reverse complement sequences were added into the *path* domain of the P strand, and further NUPACK simulations were conducted under experimental salt conditions (50 mM $Na^{+}$, 20 mM $Mg^{2+}$) to rule out undesired interactions. In the example of SARS-CoV-2 target site, sequence 2 (Figure SN2a) yielded the most favorable predicted outcome and was thus selected for experimental validation of the method.

To achieve a high diagnostic performance, increasing the *efficiency*, that is, the percentage of triggers leading to the desired strand displacement output under equilibrium conditions, was important. Similar theoretical metrics such as *availability* have been proposed previously to evaluate strand displacement reactions[6]. This importance of TMSD efficiency is apparent in main text Figure 3a (lane 4) where, despite thermal annealing, the constructs do not lead to quantitative formation of the desired product.

Consequently, the annealing assay represents the fraction of constructs available for strand displacement which could be preliminarily forecasted using NUPACK analysis. To recreate this computationally, we tested the strand displacement product formation at varying conditions, such as different concentrations (10 nM - 1000 nM), amounts of blocking strands (1- 20 eq.), and temperatures (4 °C, 25 °C, and 37 °C). For this, we evaluated the efficiency of the TMSD cascade using NUPACK both in the presence (Figure SN2b) and absence (Figure SN2c) of pathogen sequences.

### 3 Supplementary Figures

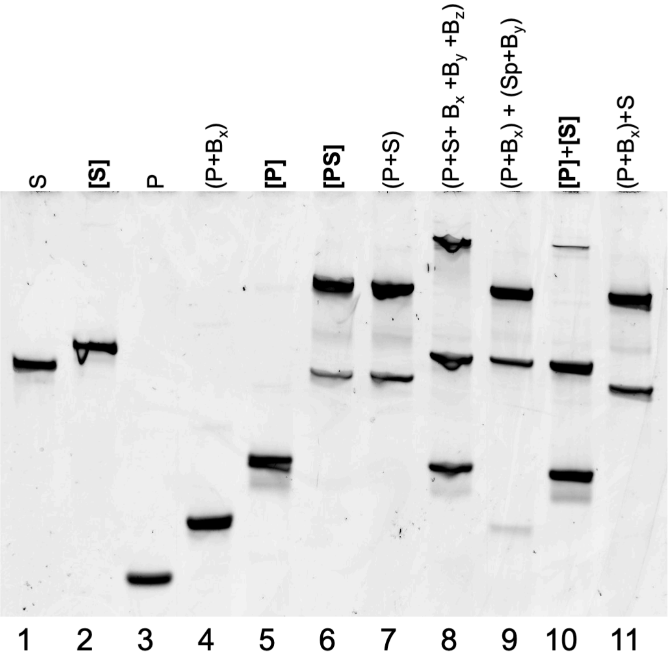


**Figure S1.** Initial validation of the TMSD cascade shows the interactions between different strands, as shown in the form of shifts in band positions. We also noted the positions of different intermediates TMSD cascades, behavior in the presence and absence of critical blocking strands, and predictions of leakage products. Square brackets indicate constructs that were pre-annealed. Round brackets indicate strand mixtures that were mixed but not annealed.

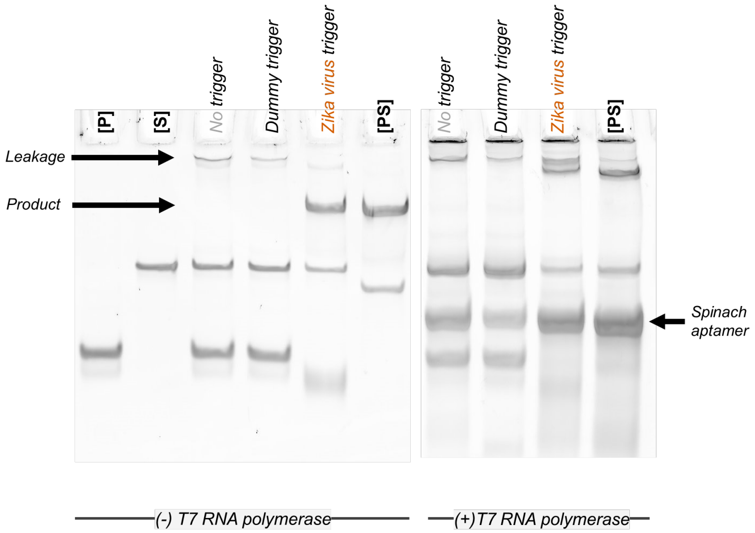


**Figure S2.** PAGE analysis shows that integrating the TMSD cascade with IVT leads to the formation of RNA aptamer. The spinach aptamer has an affinity to bind to template which can be noticed by the shift in the position of leakage/product after IVT. Integrating IVT with TMSD cascade amplifies the signal after strand displacement.

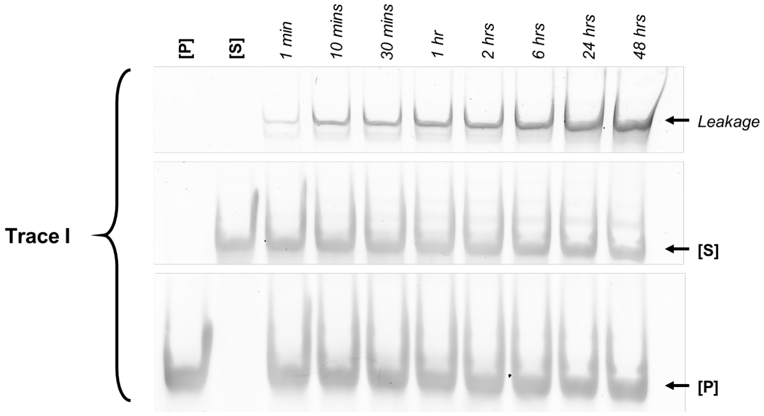


**Figure S3.** Leakage profile of Trace I showing the leakage at different time points of incubation of initial constructs. This experiment was conducted once, n=1.

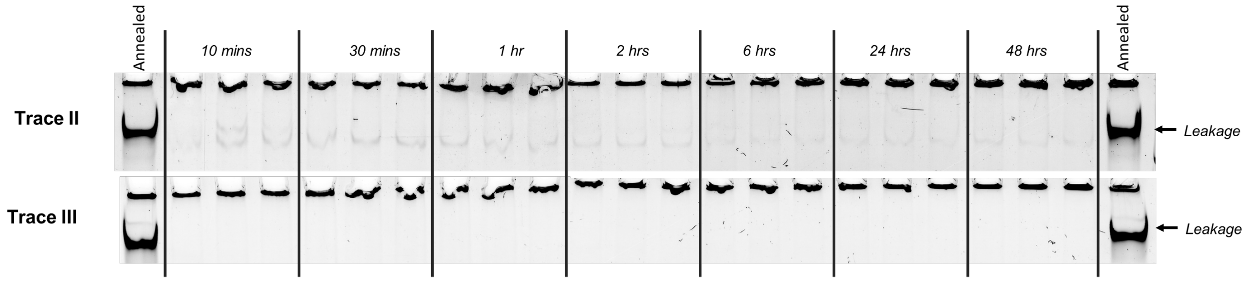


**Figure S4.** Leakage profile after employing design optimizations of leakage-reduced (Trace II) and leakage-tolerant (Trace III) cascaded at different time points of incubation of constructs. The experiment was conducted in triplicates, n=3. The dark band at the bottom of the wells is due to recombinant BSA used in the assay. Annealed samples work as reference representing 100% leakage.

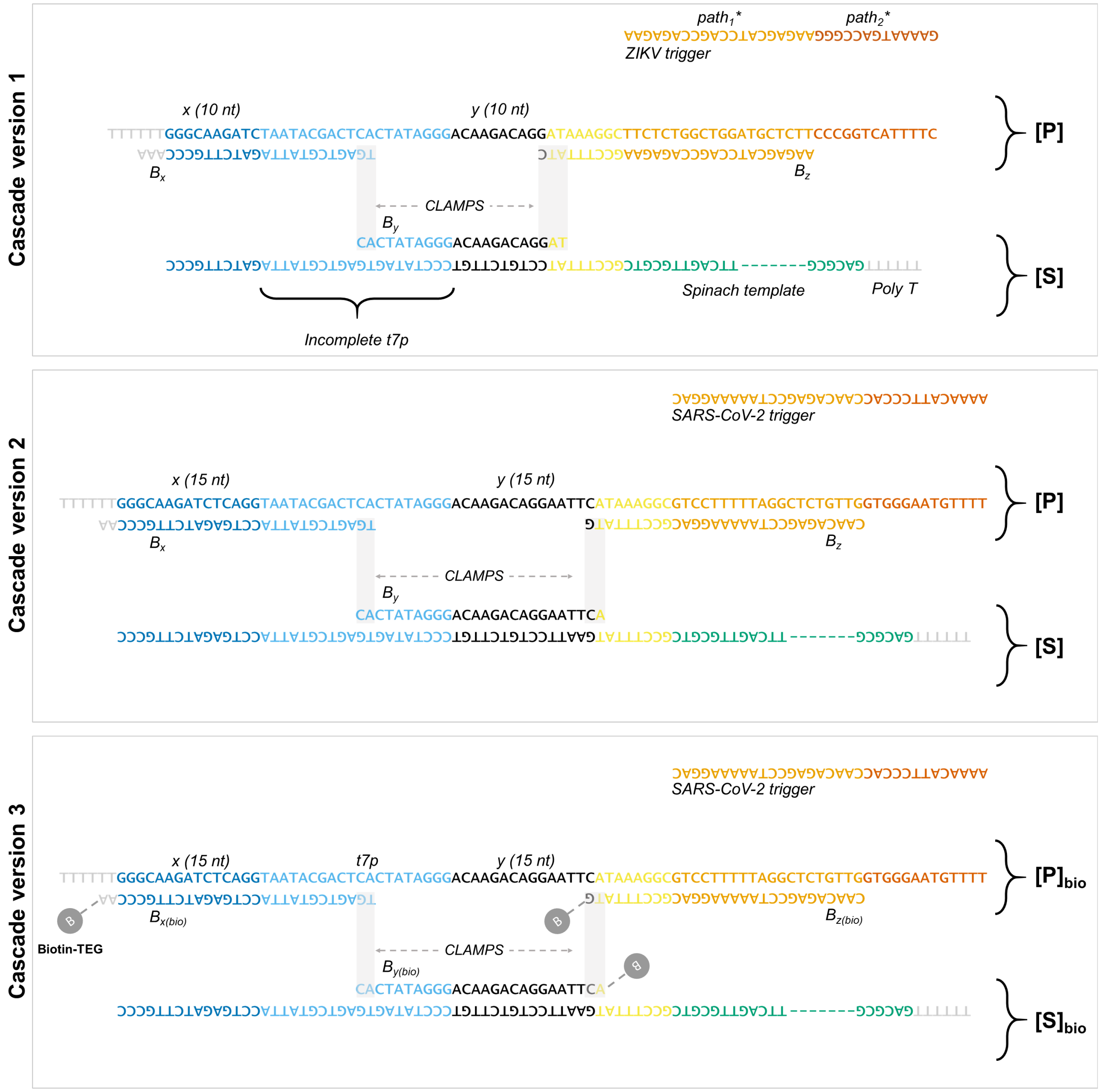


*Spinach template (98 nt) - GACGCAACTGAATGAAATGGTGAAGGACGGGTCCAGGTGTGGCTGCTTCGGCAGTGCAGCTTGTTGAGTAGAGTGTGAGCTCCGTAACTAGTCGCGTC

**Figure S5.** Aligned sequences of **[P]** and **[S]** constructs in the different cascade versions (cf. main text Figure 3b).

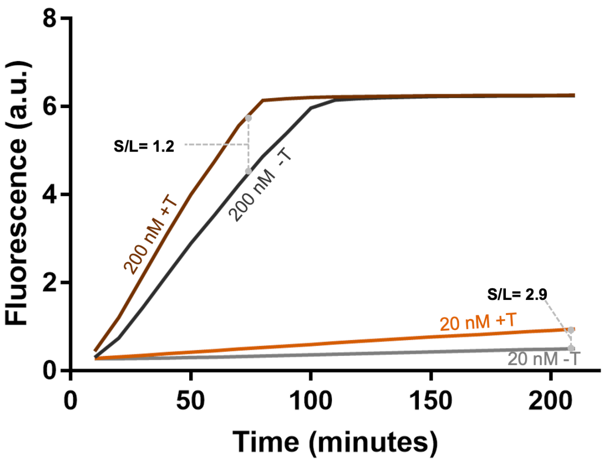


**Figure S6.** Real-time fluorescence assay shows that a reduced concentration of **[P]** and **[S]** constructs leads to a notably higher signal-to-leakage ratio.

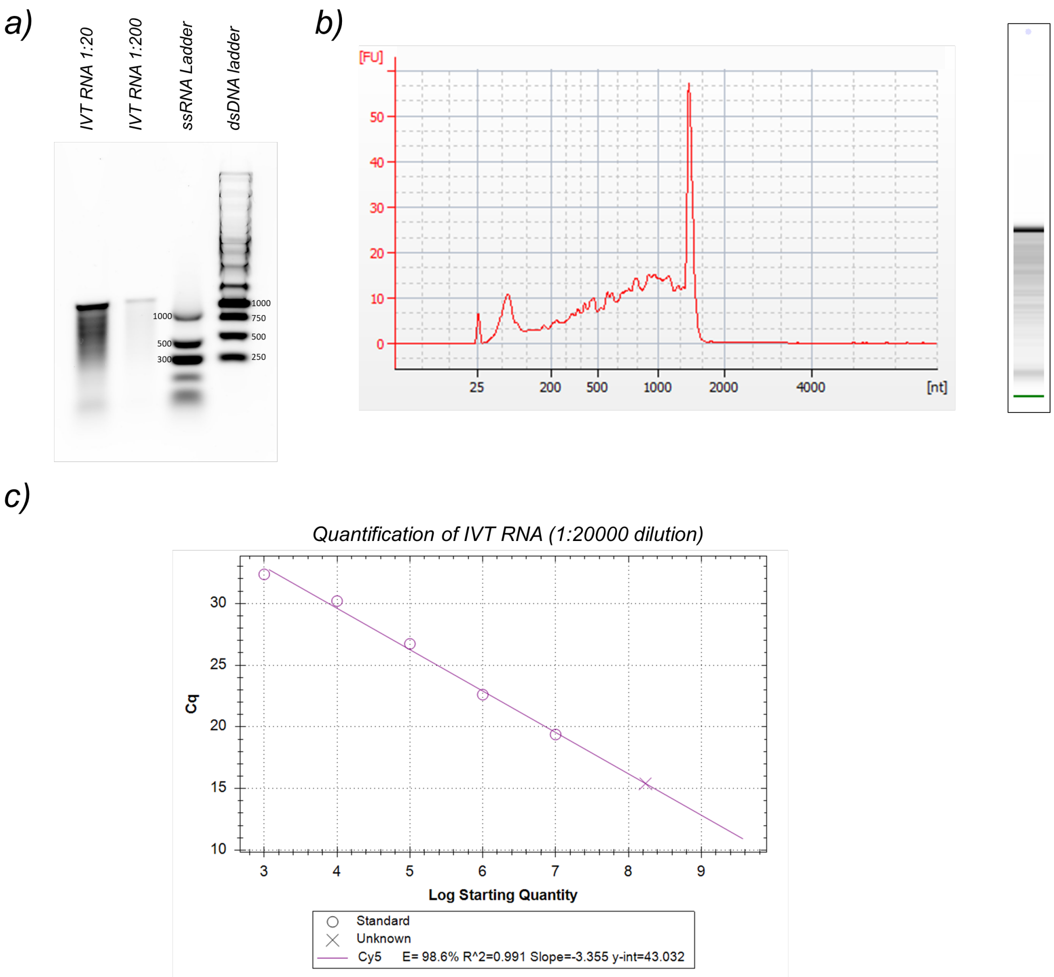


**Figure S7.** In vitro transcription, validation, and quantification of N gene SARS-CoV-2 mimic. a) Different concentrations of ~1.3 kb in vitro transcribed RNA on a 1.5% agarose gel. b) Plots from Bioanalyzer 2100 instrument. The electrophoresis lane is also shown on the side c) Typical standard curves for the quantification of SARS-CoV-2 RNA using RT-qPCR. The x-axis indicates the number of copies of the RNA reference (B.1.1.7) per tube.

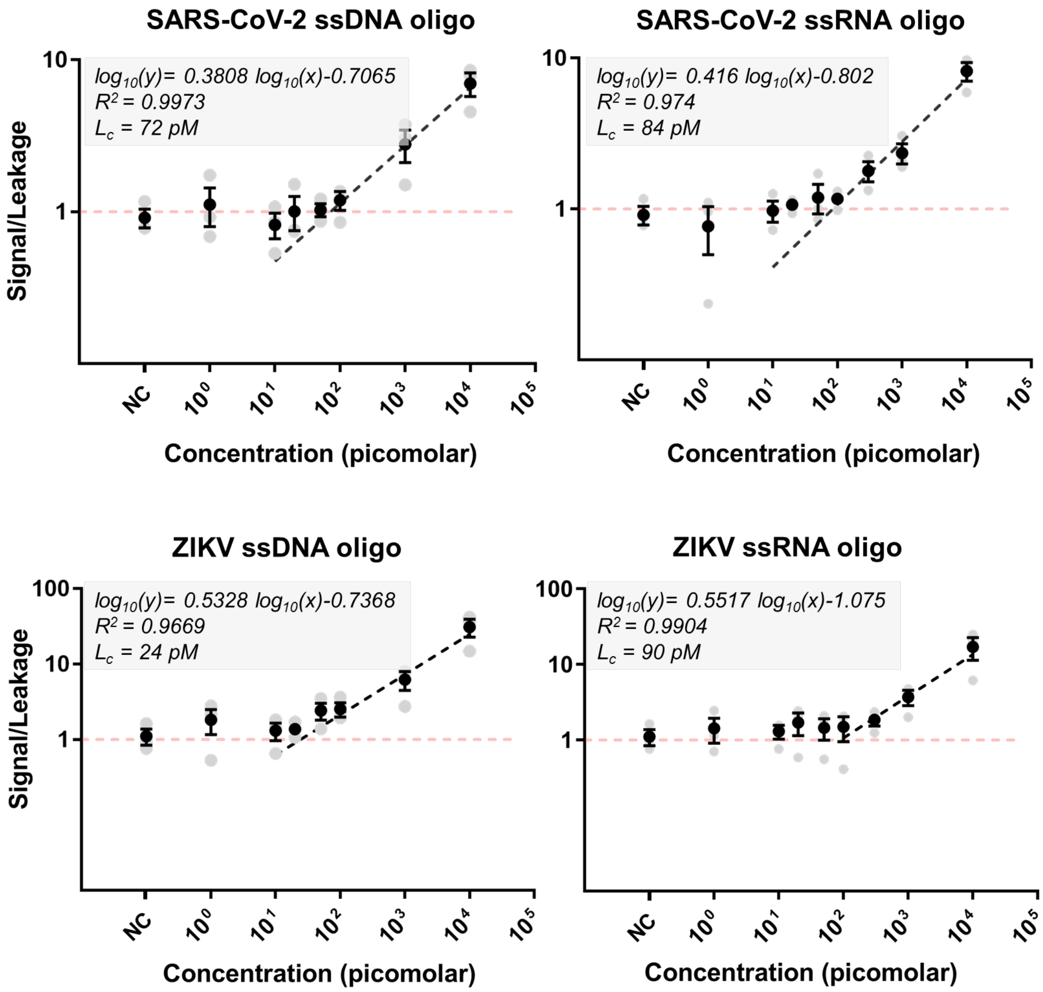


**Figure S8.** Sensitivity plots of the SARS-CoV-2-and ZIKV-specific detection assays using different concentrations of 33-nt long DNA and RNA targets after design improvements. $L_c$ is estimated using linear regression and provides a robust estimate of the theoretical detection limit. It is determined as the intersection of the linear regression of $slope_{signal}/slope_{leakage}$ (S/L) with the threshold of 1. NC = negative control containing cytoplasmic RNA. The data was collected from three independent experiments (n=3).

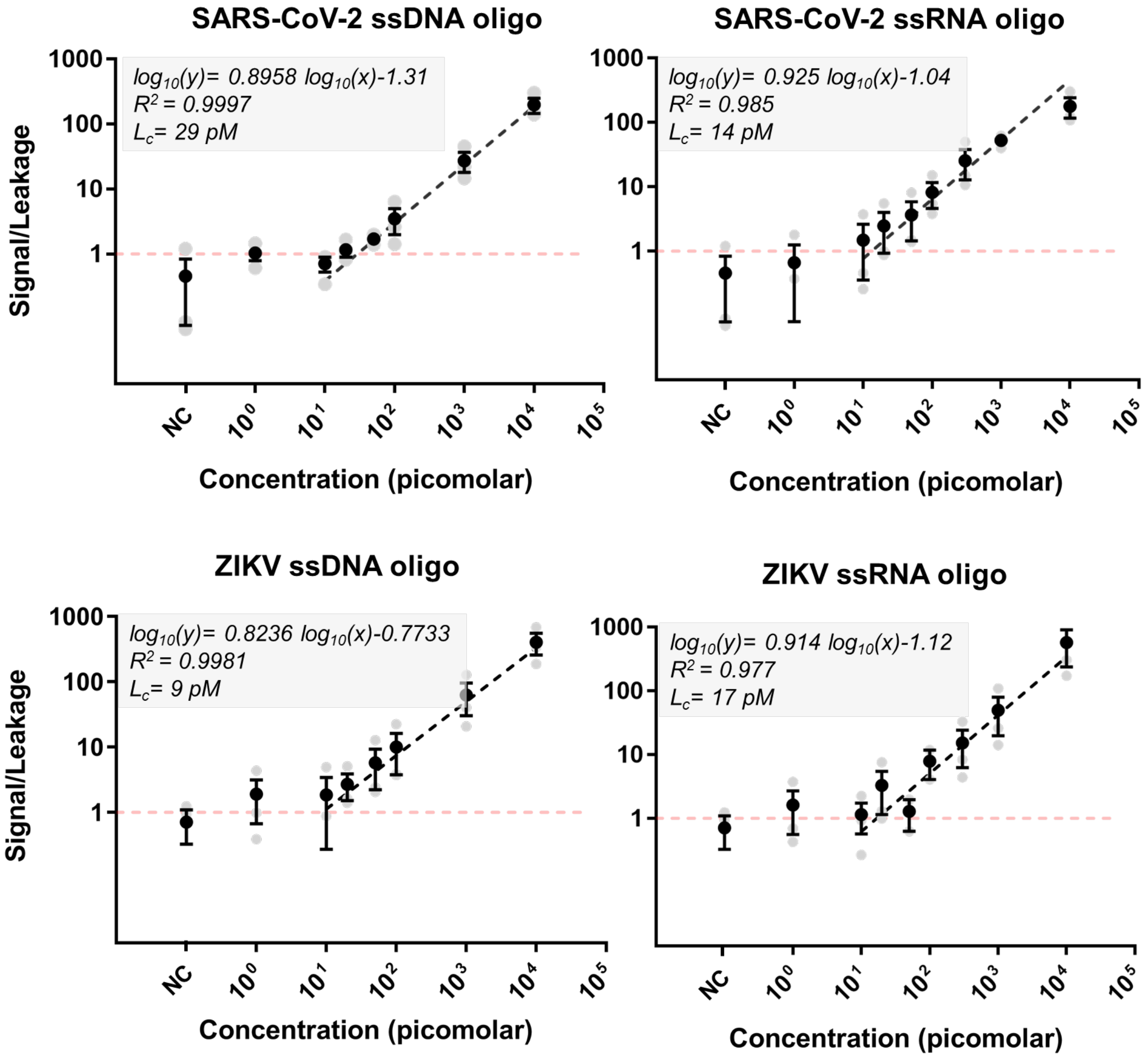


**Figure S9.** Sensitivity plots of the leakage-tolerant SARS-CoV-2-and ZIKV-specific detection assays using different concentrations of 33-nt long DNA and RNA targets. The S/L ratio improves by >10 times. This leads to comparable values of $L_c$ and LoD, improving the theoretical estimation of the analytical detection limit offered by the assay. LoD is estimated as the intersection of linear regression of $slope_{signal}/slope_{leakage}$ with the threshold of 1. NC = negative control containing cytoplasmic RNA. The data was collected from three independent experiments (n=3).

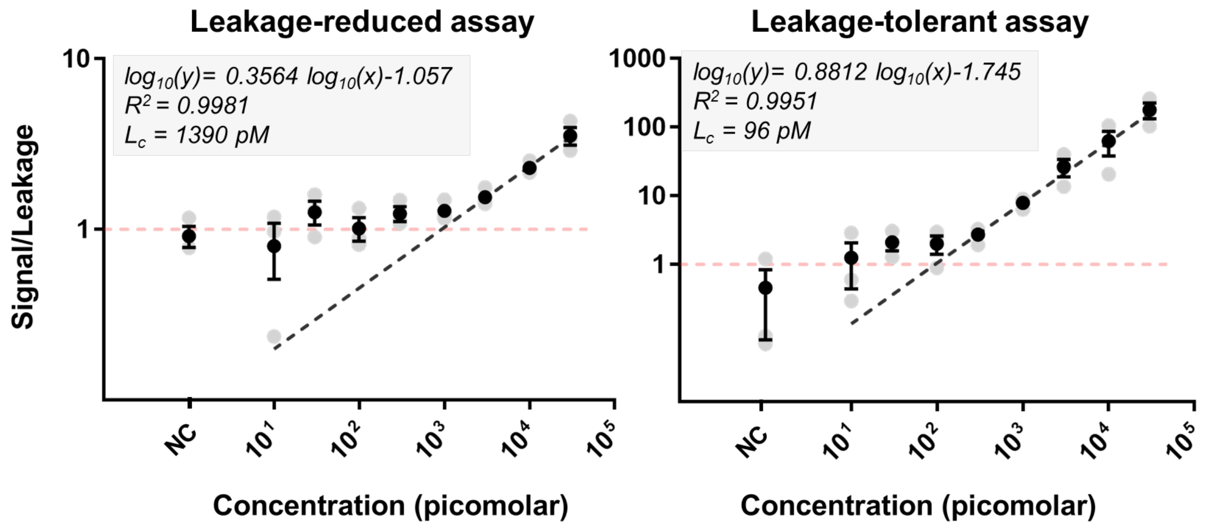


**Figure S10.** Sensitivity plots of the 1.2 kb long SARS-CoV-2 mimic using leakage-reduced and leakage-tolerant assays. The improvement in signal-to-noise ratio is evident as it improves over 10-fold. NC = negative control containing cytoplasmic RNA. The data was collected from three independent experiments (n=3).

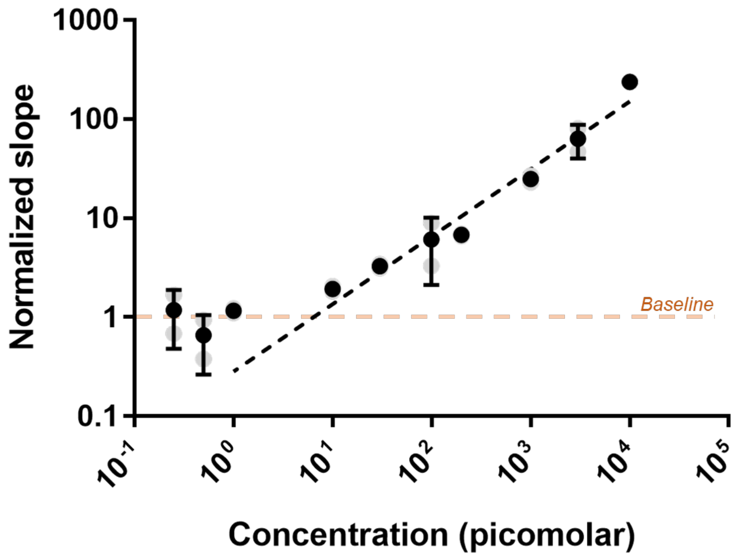


**Figure S11.** In vitro transcription of different concentrations of annealed **[PS]** template (version 2) reveals that Spinach expression cannot be detected under a concentration of ~8 pM of the activated construct. This value simulates the (theoretical) $L_c$ value for a flawlessly functioning TMSD cascade. The data was collected from two independent repeat experiments.

### 4 Supplementary Tables

**Table S1.** Predicted binding energy (ΔG) for SARS-CoV-2 and ZKV detection constructs after NUPACK design. The NUPACK simulations were performed at a concentration of 20 nM for the constructs, 10 equivalents (200 nM) of blocking strands, and 1 equivalent (20 nM) of trigger. The salt concentrations used were 50 mM $Na^+$ and 20 mM $Mg^{2+}$. These values are for the optimized Zika virus and SARS-CoV-2 constructs with 15 nt-long x and y domains.

| Target | Construct | ΔG (kJ $mol^{-1}$) |
|---|---|---|
| SARS-CoV-2 | **[P]** construct | -309.53 |
| | **[S]** construct | -185.64 |
| | Leakage product **[PS]** | -468.57 |
| | Trigger-induced strand displacement product | -527.48 |
| Zika virus | **[P]** construct | -279.57 |
| | **[S]** construct | -163.68 |
| | Leakage product **[PS]** | -420.28 |
| | Trigger-induced strand displacement product | -487.73 |

**Table S2.** List of the DNA sequences used in this study.

| Strand | Pathogen-specific target sequences | Length [nt] |
|---|---|---|
| Dummy_ssDNA | TTTTCGCCAAGTAACTCTTTCCGGTTGTTTGCCCGCCAGATTAGAGCCTTATGA GCCGTC | 60 |
| ZIKV_ssDNA | GAAAATGACCGGGAAGAGCATCCAGCCAGAGAA | 33 |
| ZIKV_ssRNA | GAAAAUGACCGGGAAGAGCAUCCAGCCAGAGAA | 33 |
| COVID_ssDNA | AAAACATTCCCACCAACAGAGCCTAAAAAGGAC | 33 |
| COVID_ssRNA | AAAACAUUCCCACCAACAGAGCCUAAAAAGGAC | 33 |
| **Strand** | **TMSD Cascade 0 (Zika virus target)** | **Length [nt]** |
| P0 | TTTTTTGGGCAAGATCTAATACGACTCACTATAGGGACAAGACAGGATAAAGGC AATGGGAAGGAAAGAAGAGG | 74 |
| S0 | TTTTTTGACGCGACTAGTTACGGAGCTCACACTCTACTCAACAAGCTGCACTGC CGAAGCAGCCACACCTGGACCCGTCCTTCACCATTTCATTCAGTTGCGTCGCCT TTATCCTGTCTTGTCCCTATAGTGAGTCGTATTAGATCTTGCCC | 152 |
| Bx0 | TTAAGTGAGTCGTATTAGATCTTGCCCAAA | 31 |
| By0 | TTTCACTATAGGGACAAGACAGGAT | 25 |
| Bz0 | TGCCCGATAACCTCTTCTTTCCTTCCCATTGCCTTTATC | 39 |
| **Strand** | **Cascade 1_ 10nt (Zika virus target)** | **Length [nt]** |
| P1 | TTTTTTGGGCAAGATCTAATACGACTCACTATAGGGACAAGACAGGATAAAGGC TTCTCTGGCTGGATGCTCTTCCCGGTCATTTTC | 87 |
| S1 | TTTTTTGACGCGACTAGTTACGGAGCTCACACTCTACTCAACAAGCTGCACTGC CGAAGCAGCCACACCTGGACCCGTCCTTCACCATTTCATTCAGTTGCGTCGCCT TTATCCTGTCTTGTCCCTATAGTGAGTCGTATTAGATCTTGCCC | 152 |
| Bx1 | TTAAGTGAGTCGTATTAGATCTTGCCCAAA | 31 |
| By1 | TTTCACTATAGGGACAAGACAGGAT | 25 |
| Bz1 | AAGAGCATCCAGCCAGAGAAGCCTTTATC | 29 |

| Strand | Cascade 2_15nt_COVID (SARS-CoV-2 target) | Length [nt] |
|---|---|---|
| P2 | TTTTTTGGGCAAGATCTCAGGTAATACGACTCACTATAGGGACAAGACAGGAAT TCATAAAGGCGTCCTTTTTAGGCTCTGTTGGTGGGAATGTTTT | 97 |
| S2 | TTTTTTGACGCGACTAGTTACGGAGCTCACACTCTACTCAACAAGCTGCACTGC CGAAGCAGCCACACCTGGACCCGTCCTTCACCATTTCATTCAGTTGCGTCGCCT TTATGAATTCCTGTCTTGTCCCTATAGTGAGTCGTATTACCTGAGATCTTGCCC | 162 |
| Bx2 | TGAGTCGTATTACCTGAGATCTTGCCCAA | 29 |
| By2 | CACTATAGGGACAAGACAGGAATTCA | 26 |
| Bz2 | CAACAGAGCCTAAAAAGGACGCCTTTATG | 29 |
| Bx2(bio) | TGAGTCGTATTACCTGAGATCTTGCCCAA/3BioTEG/ | 29 |
| By2(bio) | CACTATAGGGACAAGACAGGAATTCA/3BioTEG/ | 26 |
| Bz2(bio) | CAACAGAGCCTAAAAAGGACGCCTTTATG/3BioTEG/ | 29 |
| Strand | Cascade 2_15nt_ZIKV (Zika virus target) | Length [nt] |
| P3 | TTTTTTGGGCAAGATCTCAGGTAATACGACTCACTATAGGGACAAGACAGGAAT TCATAAAGGCTTCTCTGGCTGGATGCTCTTCCCGGTCATTTTC | 97 |
| S3 | TTTTTTGACGCGACTAGTTACGGAGCTCACACTCTACTCAACAAGCTGCACTGC CGAAGCAGCCACACCTGGACCCGTCCTTCACCATTTCATTCAGTTGCGTCGCCT TTATGAATTCCTGTCTTGTCCCTATAGTGAGTCGTATTACCTGAGATCTTGCCC | 162 |
| Bx3 | TGAGTCGTATTACCTGAGATCTTGCCCAA | 29 |
| By3 | CACTATAGGGACAAGACAGGAATTCA | 26 |
| Bz3 | AAGAGCATCCAGCCAGAGAAGCCTTTATG | 29 |
| Bx3(bio) | TGAGTCGTATTACCTGAGATCTTGCCCAA/3BioTEG/ | 29 |
| By3(bio) | CACTATAGGGACAAGACAGGAATTCA/3BioTEG/ | 26 |
| Bz3(bio) | AAGAGCATCCAGCCAGAGAAGCCTTTATG/3BioTEG/ | 29 |

/3BioTEG/ = 3' Biotin-TEG modification

**Table S3.** Primers used for amplification of N gene DNA template with T7 adapter.

| Strand | SARS-CoV-2 transcription template | Length [nt] |
|---|---|---|
| N_Gene_FW_T7 | TAATACGACTCACTATAGGGCGATGACGATGGATAGCGA | 39 |
| N_Gene_REV | TGTGTGGAATTGTGAGCGGA | 20 |

Red sequence = T7 adapter

**Table S4.** qPCR program used for the amplification of N gene PCR product.

| Temp | Time | Cycles |
|---|---|---|
| 98 °C | 1 minute | |
| 98 °C<br>68 °C<br>72 °C (Plate Read) | 15 seconds<br>15 seconds<br>60 seconds | 30 cycles |
| 72 °C | 3 mins | |
| 55- 95 °C | | Melt curve |

**Table S5.** Calculation of improvements in leakage due to design improvements.

| | Leakage (%) 2 hr | Leakage (%) 48 hr | Slope (%/hr) | Inherent leak reduction (%) |
|---|---|---|---|---|
| Trace I | 20.14 | 41.83 | 0.47 | 96.58 |
| Trace II | 2.00 | 2.75 | 0.02 | |

| | Leakage (%) Trace I | Leakage (%) Trace II | Defect Leakage reduction (%) |
|---|---|---|---|
| 1 hr | 18.70 | 1.50 | 91.96 |

**Table S6.** Calculations for the efficiency of leakage capture using streptavidin-coated magnetic beads. The quantification was conducted twice and values were averaged to reduce calculation errors.

| | Quantification 1 | | | Quantification 2 | | |
|---|---|---|---|---|---|---|
| | Leakage | [S] | [P] | Leakage | [S] | [P] |
| before capture | 10263.49 | 8312.00 | 7549.00 | 11465.08 | 10640.32 | 9334.83 |
| after capture | 379.85 | 290.00 | 165.00 | 376.09 | 256.85 | 208.44 |
| efficiency (%) | 96.30 | 96.51 | 97.81 | 96.72 | 97.59 | 97.77 |

| Average capture efficiency (%) | |
|---|---|
| [Leakage] | 96.51 |
| [S] | 97.05 |
| [P] | 97.79 |
